# Conformational dynamics of active site loops 5, 6 and 7 of enzyme Triosephosphate Isomerase: A molecular dynamics study

**DOI:** 10.1101/459198

**Authors:** Sarath Chandra Dantu, Gerrit Groenhof

## Abstract

Triosephosphate Isomerase is a glycolytic enzyme catalyzing the interconversion of Dihydroxyacetone phosphate to Glyceraldehyde-3-phosphate. The active site is comprised of three distinct loops loop-6, loop-7 and loop-8. Based on loop-6 and loop-7 conformation we describe the enzyme as Open TIM and Closed TIM. Various NMR, X-ray crystallography and QM/MM simulation techniques have provided glimpses of individual events of what is essentially a dynamic process. We studied the conformational changes of two distinct loops (loop-6 and loop-7) enveloping the active site, in the presence of natural substrate, reaction intermediates and inhibitor molecules, by means of microsecond atomistic MD simulations in solution and crystal environment. Our studies have revealed that loop-6 samples open and closed conformations in both apo and holo TIM structures. As seen in solution state NMR experiments, we also observe that loop-6 N-terminus and C-terminus move independently. In our simulations we have also observed that backbone dihedrals of loop-7 residues G210 (G210-phi, G210-psi) and G211 (G211-phi) sample open and closed states in both apo and holo TIM structures. Whereas backbone dihedral angles of G211 (G211-psi) and S212 (S212-phi) adopt closed conformation only when the ligand is bound to the active site. As observed in chain-B of 1R2R crystal structures, we also observe that water molecules can also initiate flip of G211-psi and S212-phi dihedral angles into closed conformation. Except, loop-5, which has a dominant effect on the conformational behaviour of loop-6 N-terminus, we do not observe any influence of either loop-6 or loop-7 on the conformational dynamics of the other.

## 1 Introduction

Glycolysis involves conversion of six carbon compound glucose, to two molecules of three carbon compound pyruvate, harvesting two molecules of Adenosine triphosphate (ATP). The fourth reaction of this metabolic pathway involves conversion of Fructose 1, 6-Bisphosphate to a ketone compound Dihydroxyacetone Phosphate (DHAP) and an aldehyde compound Glyceraldehyde-3-Phosphate (GAP or G3H). GAP proceeds further to form one pyruvate molecule, whereas DHAP is converted to GAP by TIM in the fifth step of glycolysis, which in turn forms the second pyruvate. Interconversion of DHAP to GAP and GAP to DHAP is catalyzed by TIM in both the directions^1^ at a catalytic rate (k_cat_) of 10^3^ s^−1^ and 10^4^ s^−1^ respectively.

### 1.1 TIM structure

TIM was first crystallized by Phillips et al. in 1975^2^. TIM is active as a dimer (Figure) in most of the organisms and tetramers were also reported in thermophillic organisms^1,3-5^. Mutations in the dimer interface lead to formation of monomers which exhibit a 1000 fold lower catalytic rate when compared with the wild type dimer^6^. Hence to counter diseases like malaria and African sleeping sickness various studies have concentrated on developing an inhibitor which binds at the dimer interface and disrupts the quaternary structure^7-13^ . Each monomer is composed of approximately 250 amino acids. The tertiary structure of TIM is composed of 8 α and 8 parallel β sheets interconnected by loops (**Figure 1**). This (βα)_8_ fold also known as TIM-barrel fold, is shared by more than 10% of known enzyme structures^14^.

**Figure 1:**
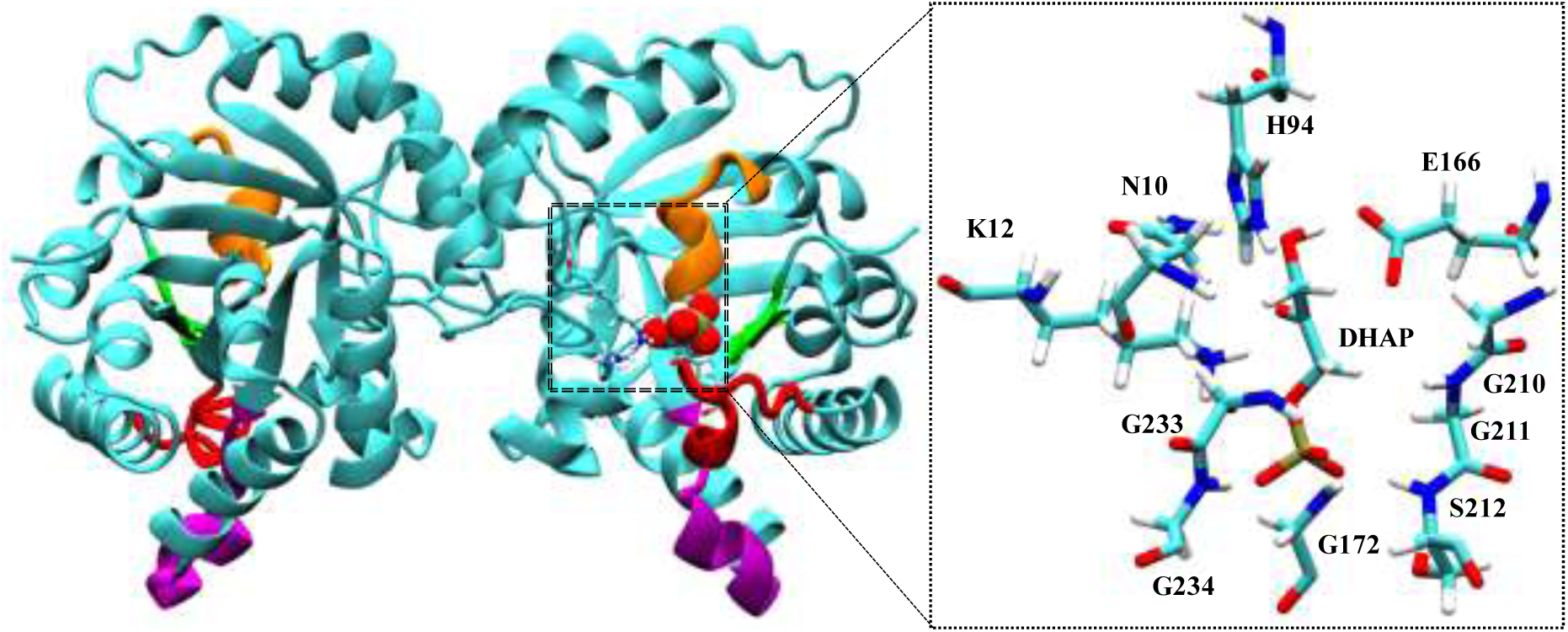
Structure of TIM dimer with DHAP in the active site. Loop-6 (red), loop-7 (green) and loop-8 (orange) envelope the active site. Loop-5 (purple) interacts with N-terminus residues of loop-6. In inset residues of the active site which are highly conserved.

The active site of TIM is enveloped by three distinct loops (**Figure 1**) loop-6, loop-7, loop-8. Residues H94, E166 are the proposed catalytic residues involved in the proton shuttling between C1 and C2 carbon atoms of the substrates. G172 (loop-6), S212 (loop-7), G233 and G234 (loop-8) provide hydrogen bonds to the phosphate moiety of the substrate and stabilize it (**Figure 1**). N10 and K12 interact electrostatically with the substrate (**Figure 1**). Residues that are involved in catalysis and that are interacting by means of hydrogen bonds or coulombic interactions are conserved throughout all the sequences^1^.

Loop-6 N-terminus (167-PVWA-170), tip (171-IGTG-174) residues are much more conserved than C-terminus (175-KVA-177) residues (**Figure 2**). Wang et al. reported that there is an evolutionary link between the sequence of N-terminus segment loop-6 (PVW) and loop-7. The typical loop-6 N-terminus sequence is ‘167PVW’ with a corresponding loop-7 sequence ‘209YGGS’. But in archaebacteria loop-6 N-terminus sequence is ‘167PPE’ and typical loop-7 sequence being ‘209TGAG/209CGAG’. A substitution of archaebacterial loop-7 sequence into chicken TIM, with unmodified loop-6, leads to an impaired enzymatic activity ^15^.

**Figure 2:**
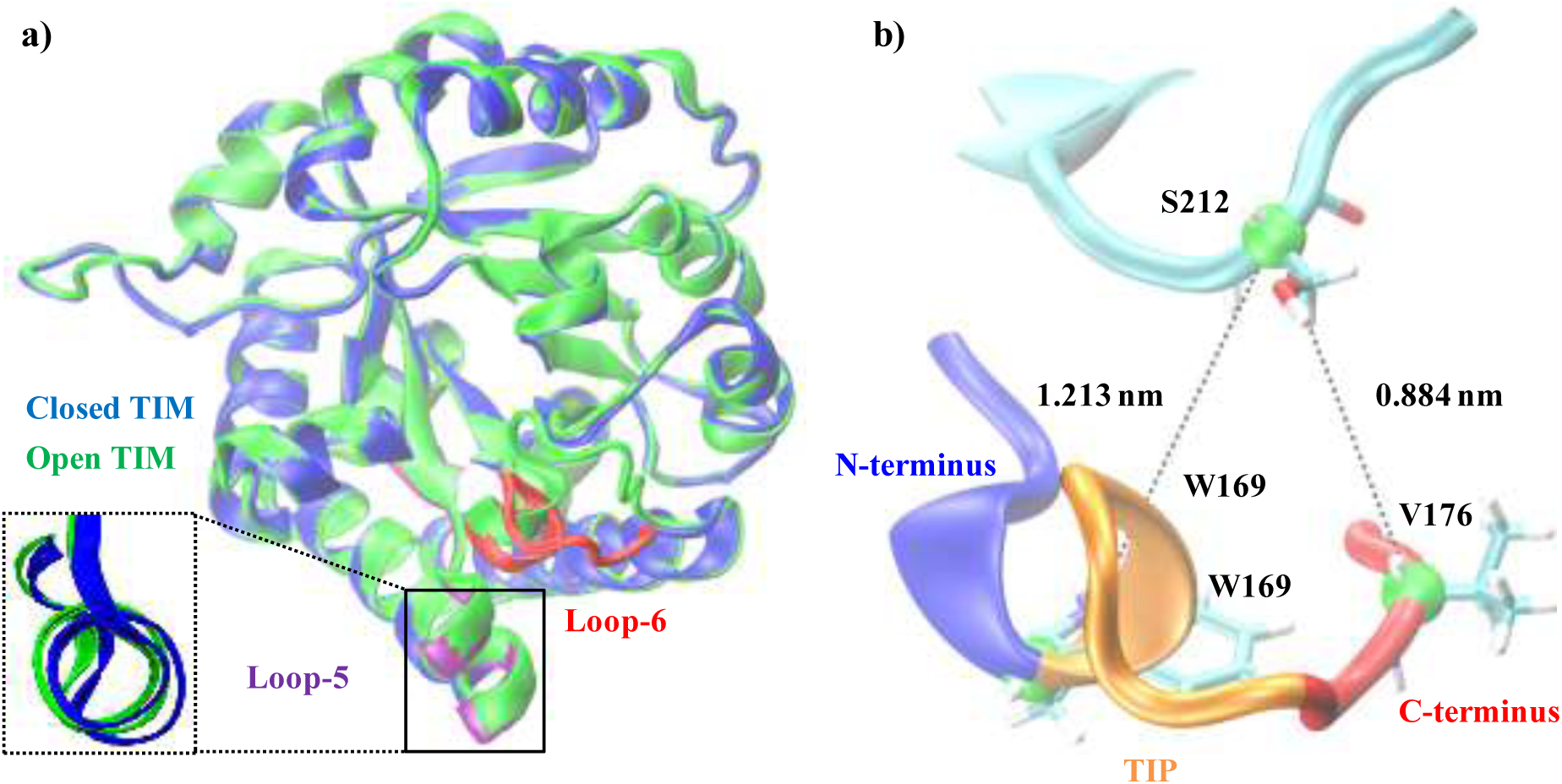
a) Fit of OTIM (green) structure on CTIM (blue). Major structural difference is seen at loop-6 (open loop-6 in red). In inset open (green) and closed (blue) loop-5. b) N-terminus, C-terminus and tip of open loop-6. In our simulations, to monitor hinge regions dynamics W169-S212 and V176-S212 c-alpha (green spheres) distances are monitored.

There are three structural differences between open TIM (OTIM: 5TIM-A) and closed TIM (CTIM: 6TIM-B) structures (**Figure 2a**). The major structural difference between OTIM and CTIM is seen in loop-6. Backbone RMSD of 5TIM-A with respect to 6TIM-B excluding loop-6 is 0.034nm. The second structural difference is seen in loop-7 where three residues undergo a change in Φ, Ψ dihedral angles between OTIM and CTIM (**Figure 2**). There is also a minor structural change seen in loop-5 between OTIM and CTIM (**Figure 2a**). Carboxylate moiety of E128 from loop-5 forms a hydrogen bond with epsilon hydrogen of loop-6 W169 in closed state (**Figure 2a**). This hydrogen bond is absent in open loop-6. Excluding loop-6 and loop-7, rest of the protein is highly identical between open and CTIM. The role of each loop-6, loop-7 and residues involved in the enzyme catalysis are further discussed below.

### 1.2 Loop-6

Loop-6 is composed of 11 amino acids (Consensus sequence: “PVWAIGTGKTA”). N-terminus of loop-6 (‘PVW’) is rich in hydrophobic amino acids whereas tip (“AIGTG”) of loop-6 and C-terminus (“KTA”) are hydrophilic. N-terminus of loop-6 interacts with the loop-5 whereas tip of the loop-6 and C-terminus are solvent exposed. Loop-6 acts as a lid on the active site. Depending upon the orientation of loop-6 upon the active site, TIM has two distinct conformations; open and closed (**Figure 3a**). We have used distance between residues W169-S212 and V176-S212 as a criterion to distinguish between open and CTIM (**Figure 3**).

**Figure 3:**
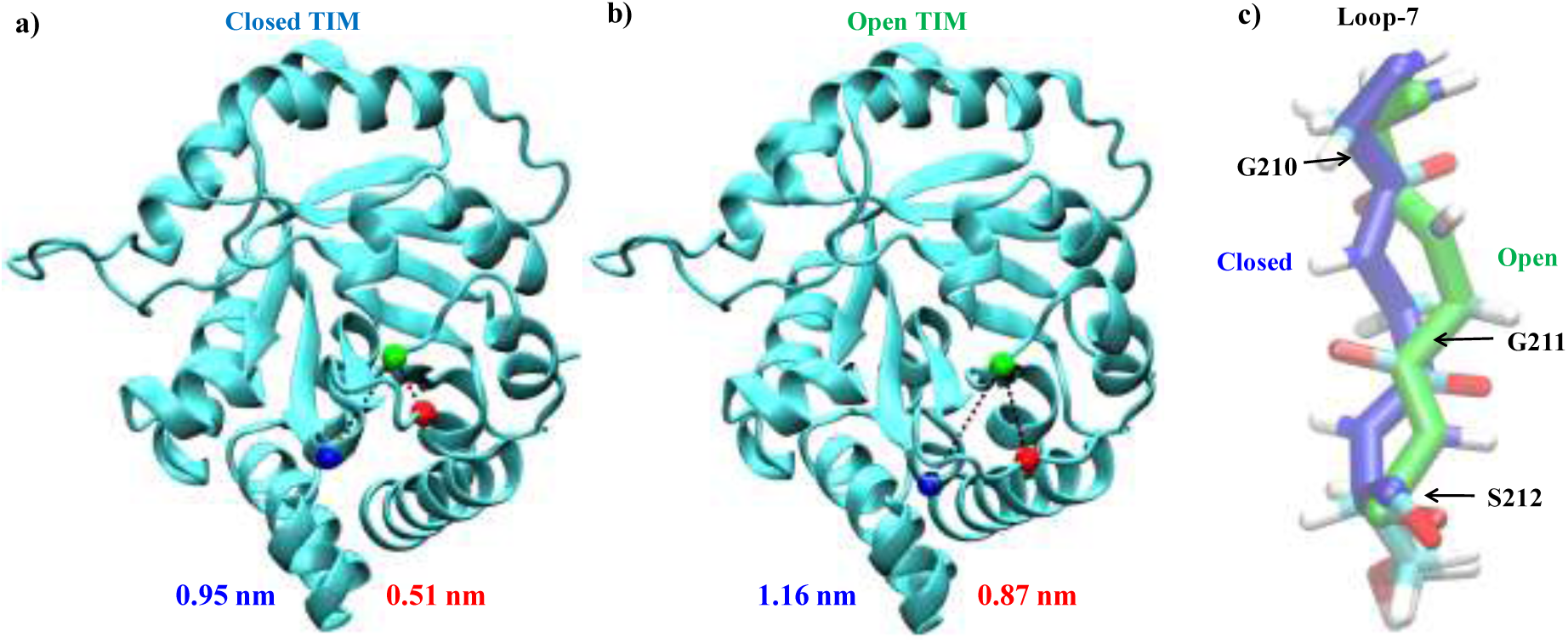
Closed (a) and open (b) conformations of TIM based on distance between W169-S212 (Closed: 0.95nm, Open: 1.16nm) and V176-S212 (Closed: 0.51nm, Open: 0.87nm). Distance to loop-6 (red) and loop-7(red). c) Loop-7 in open (green) and closed conformation (blue).

A mutation study by Pompliano et al. where loop-6 residues ‘171IGTG’ were deleted, suggested that closure of loop-6 during the enzyme catalysis is assumed to stabilize the enediol phosphate intermediate as well as the transition state^16^. In the mutant TIM they found a six fold higher formation of methylglyoxal and inorganic phosphate compared to the wild type enzyme^16^. The so formed methylglyoxal is toxic to the cell^17^.

#### 1.2.1 NMR studies on loop-6

As stated earlier, dynamic nature of loop-6 has been investigated by several scientific studies. W169 present in the N-terminus of loop-6 was used as an indicator of the loop-6 conformation in the studies conducted by Ann McDermott group^18-21^. There is change in the environment of the W169 between the two conformations. In open loop-6, epsilon hydrogen of W169 is involved in a hydrogen bond with hydroxyl group of Y165. In closed loop-6, epsilon hydrogen of W169 forms hydrogen bond with carboxylate moiety of E128 (**Figure 4**). Upon closure of loop-6 W169 swings by 30°-40° (**Figure 4c**). Patrick Loria group^15,22^ in their solution NMR relaxation dispersion experiments monitored the dynamics of loop-6 N-terminus V168 and C-terminus K175, T178 residues.

**Figure 4:**
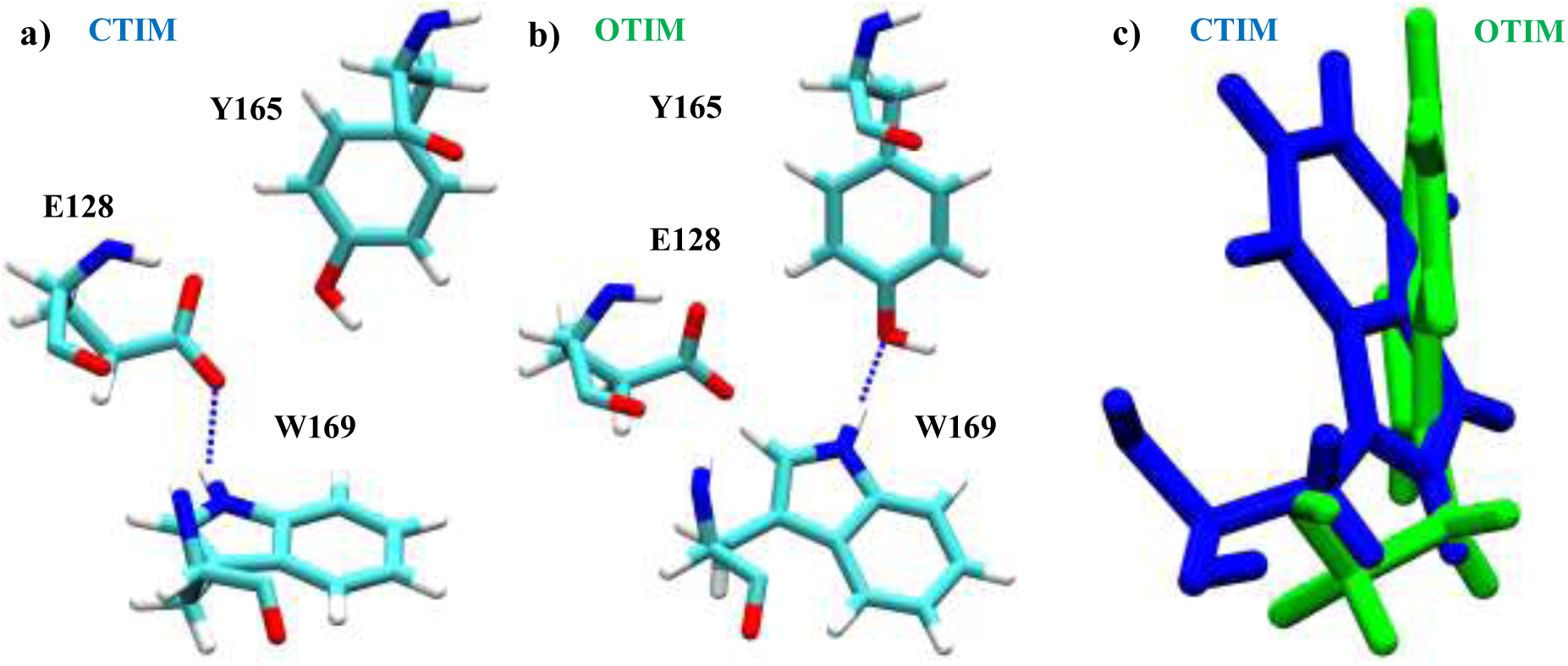
a, b) W169 forms hydrogen bond with E128 in CTIM and with Y165 in OTIM.

Solid state NMR (SSNMR) study by Williams et al. in 1995 of apo and Glycerol-3-Phosphate (G3P) ligated TIM reported that loop-6 motion is not ligand gated and occurs at a rate of 10^4^s^−1^ . In SSNMR the obtained deuterium line shapes are simulated with ligand concentration, W169 angle and loop-6 populations as parameters. Based on the crystallographic data Williams et al. assumed that in their NMR experiments, in apo state open loop-6 is the major conformation and in the ligated TIM closed loop-6 is the major conformation. Using the exchange rates and population ratios obtained from the simulated line shapes, barrier heights and free energy difference between open and closed conformations of apo TIM and G3P/PGA ligated TIM was calculated. The reported free energy difference between the major conformation and the minor conformation ranges between 1.2 kcal.mol^−1^ and 1.8 kcal.mol^−1^ . If the binding free energies of G3P and 2PG molecules are also considered along with the difference in free energy between open and CTIM’s, then it would be expected that in apo state, OTIM is favoured by −1.8 kcal.mol^−1^ and in ligated TIM with G3P/2PG closed state is favoured by −4.5/−6.8 kcal.mol^−1^ with respect to apo OTIM^18^.

T-jump relaxation studies by Desamero et.al^21^ in 2002 and solid, solution state NMR studies by Rozovsky et al.^19^ in 2003, reported that loop-6 opening rate in presence of G3P, is of the same order as the catalytic rate of TIM which is 10^4^s^−1^. Desamero et.al^21^ further suggested that loop-6 opening rate depends upon the ligand present in the active site. It has to be noted that the opening and closing rates reported by the above studies are the k_close_ and k_open_ rates (**Supplementary Figure 1**). It was difficult to measure the k_ex_ in apo TIM sample using fluorinated W168 in solution state NMR studies because no change in chemical shift is observed and Rozovsky et al.^19^ suggested that it might because of low population of closed loop-6. It is speculated though that in apo state loop-6 motion is in the regime of 10^5^s^−1^ − 10^6^s^−1^ ^20^.k_on_ and k_off_ rates were proposed to be 10^6^s^−1^.^20^

By mutational studies and using TROSY NMR spin relaxation experiments, Berlow et al. ^22^ has shown that N and C-terminal hinges of loop-6 are coupled. Y209F mutation in loop-7 destabilizes the closed conformation of loop-6 and increases the population of open state because of the loss of hydrogen bond between C-terminus loop-6 residue A176 and Y209. The effect of this mutation is seen in the dynamics of both N and C-terminus hinges. Surprisingly they found different populations distributions for N and C-terminal hinges, which might imply that N and C-terminus hinges of loop-6 are uncoupled and they suggest that more experiments are required to confirm this “novel” view of loop-6 motion^22^.

Further NMR studies by Wang et al.^15^ where chicken loop-7 (208YGGS) was replaced by archaeal loop-7 sequence (208TGAG), reported a 240 fold loss of activity for TIM. The affect of this loop-7 mutation on the exchange rate (k_ex_) between open and closed conformations of loop-6 for V167 and T177 residues was identical (wild type: 9000s^−1^; loop-7 mutant: 18000s^−1^) and based on this data they suggested that in the absence of the proper corresponding loop-7 sequence, concerted motion between the N and C-terminal hinges of loop-6 is affected. Wang et al. ^15^ also calculated the height of the activation barriers for loop-6 closure for V168 and T177 in wild type enzyme (V167: 13.7±1.8 kcal.mol^−1^; T177: 15.7±0.98 kcal.mol^−1^) and as well as the loop-7 mutant (V167: 1.4±0.38 kcal.mol^−1^; T177: 6.26±2.6 kcal.mol^−1^)^15^ .

The barrier heights are calculated by measuring the change in contribution of chemical exchange process to the relaxation rate which is R_ex_ by varying temperature. Though the effect on the obtained k_ex_ rate in the loop-7 mutant was identical (k_ex_ wild type: 9000 s^−1^ , loop-7 mutant: 18,000 s^−1^) on N and C-terminus hinges, the height of the activation barriers was different and Wang et al.^15^ pointed out that this difference may be either because of difference in population rations for N and C-terminus hinges in open and closed states or the effect of temperature on the population of V167 and T177 in open and closed states is different in the mutant^15^.

### 1.3 Loop-7

Loop-7 (**Figure 5a**) is located above the active site and is composed of six amino acids (‘YGGSVN’). Phi (Φ) and psi (Ψ) backbone dihedral angles of loop-7 residues G210, G211 and S212 differ between open and CTIM structures (**Figure 5**). It was suggested that the flip of G210 (, Ψ) G211 into closed conformation induces conformational change of E166 from “swung-out” to “swung-in” reactive conformation^1^. S212 amide group is involved in a hydrogen bond formation with the phosphate moiety of the bound substrate.

**Figure 5:**
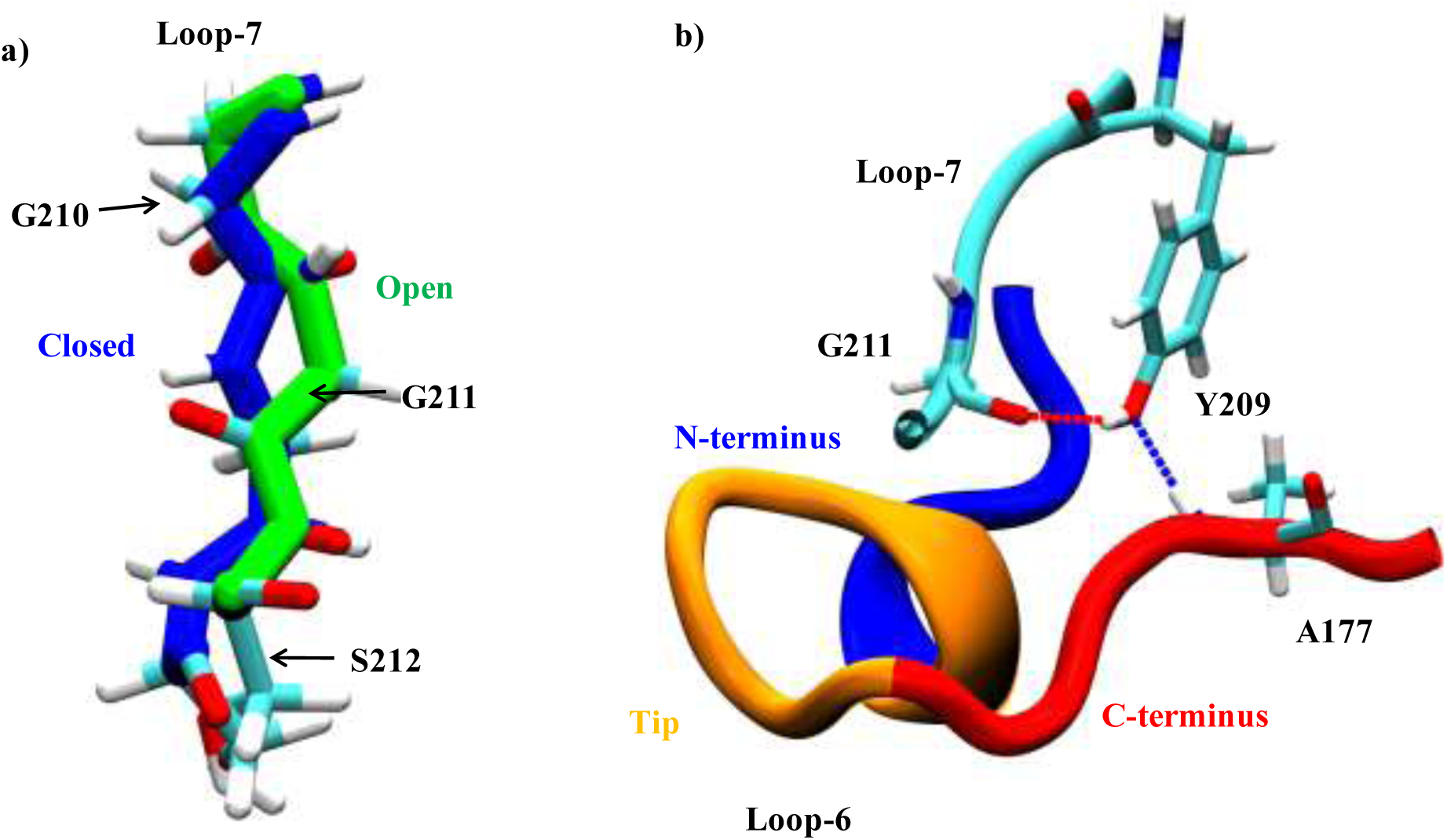
a) Loop-7 open (green) and closed (blue) conformations. b) Hydrogen bonds formed by Y209 with G211 of loop-7 and A177 of loop-6 in closed state.

Y209 of loop-7 is involved in two hydrogen bonds, one with A177 of loop-6 (**Figure 5b**) stabilizing the closed state of loop-6. The second hydrogen bond is formed between the hydroxyl group of Y209 and carbonyl oxygen of G211 (**Figure 5b**). This interaction stabilizes the G211 of loop-7 in the closed state^23^. Mutagenesis^24,25^ and NMR studies^22^ of Y209F mutant suggested that loss of hydrogen bond between the Y209 and A177 results in an increased population of the open state with a 2000 fold decrease in the catalytic activity. Loop-7 as stated above in section 1.2 was suggested to bring about concerted motions between N and C-terminus hinges.

There is a widespread interest in designing drugs to inhibit TIM, to counter malaria and African sleeping sickness^7-13^ and also designing novel enzymes based on TIM’s scaffold ^6,26-28^. Prior to that, it is important to understand step by step process of the conformational changes of loop-6 and loop-7 and the factors that drive these conformational changes. NMR and fluorescence spectroscopic studies on loop-6 have revealed insight into the exchange rates (k_ex_), between encounter complex OTIM and ligated (G3P/2PG) CTIM. The k_ex_ for apo TIM between open and closed conformations is difficult to measure by experiments because of skewed populations of loop-6 in closed conformation^19^. Previous computational studies did not simulate the entire protein and spanned only 100ns, which is not enough to capture a complete picture of such a slow (microsecond to millisecond) conformational change process.

Atomistic microsecond MD simulations of entire protein with and without ligand will help us to identify various conformational changes in TIM at atomic detail. Our study is aimed at addressing the following questions on TIM.

- What is the nature of dynamics of loop-6? Is it a rigid body motion or does N and C-terminus hinges of loop-6 move in uncorrelated fashion?
- What factors contribute to motion of loop-6? Is closed conformation sampled in apo TIM? Or is the closed conformation of loop-6 ligand induced?
- How does loop-7 influence dynamics of loop-6? Does it really bring about a concerted motion of loop-6?
- What is the nature of loop-7 conformational change? Influence of loop-7 dynamics on loop-6 has been studied but process of loop-7 conformational was not studied till now.
- Are there any other factors that influence the loop-6 and loop-7 conformational change?
- What is the free energy difference (ΔG) between open and closed conformations of loop-6 and loop-7 and what is the effect of various ligands on the ΔG?
- TIM catalyzes the reaction in both the directions i.e. binding of GAP to active site will form DHAP and vice versa. And an important question here is how does the product formed at the end of the reaction released, since the product formed at the end of previous reaction is the substrate for the next reaction?
- What changes happen in the active site upon the binding of ligand?

## 2 Methodology

To study the structure-dynamics-function relationship of TIM, we performed apo and holo TIM molecular dynamics (MD) simulations using GROMACS MD package^29-33^. We used the amber99sb^34^ all atom forcefield a variant of the Amber-99^35^ potential in which charges were derived using Quantum mechanics (QM) calculations with HF-6/31G* basis set with improved backbone chain torsion potentials. We used the AMBER ports^36^ for implementation of amber99sb in GROMACS. We have used the ion type definitions from the Joung et al.^37^.

Ligated TIM simulations were performed with natural substrate Dihydroxyacetone phosphate (DHAP), inhibitor 2-phosphoglycerate (2PG), reaction intermediates eneldioate-1 (EDT1) and eneldioate-2 (EDT2; **Supplementary Figure 2**)^38^. To keep the forcefield parameters of ligand molecules compatible with the protein forcefield, General amber forcefield (GAFF)^39^ parameters were developed for the ligand molecules which is described in section 2.4.

### 2.1 Apo TIM simulations

Apo TIM simulations were performed using 5TIM^23^ (1.83 Å), 6TIM^40^ (2.20 Å), 1NEY^41^ (1.20 Å) and 1R2R^42^ (1.50 Å) crystal structures. Structures for apo TIM simulations were created by removing the ligand molecules present in the active site. In general, all the dimer TIM crystal structures have one chain open and one chain closed. In 5TIM and 6TIM crystal structures there is no substrate/inhibitor bound to the active site of chain-A because of crystal contacts^23^ and hence chain-A is in the open conformation in both 5TIM and 6TIM. Chain-B of 5TIM has a sulfate ion and active site of chain-B of 6TIM has the substrate analogue Glycerol-3-phosphate (G3P). We have used distance between C-alpha atoms of W169 and S212, V176 and S212 to distinguish TIM structures into open and CTIM structures which will be discussed in section 3.1 and based on these distances chain-B of 5TIM and 6TIM are in closed state.

In 1R2R crystal there are two dimers per asymmetric unit i.e. four chains in total. Chains A, C and D are open, and chain-B is closed. 1R2R is the only crystal structure apart from the TIM monomeric variant 2VEI where one of the chains (chain-B) is in closed state with an unoccupied active site.

For MD simulations, we wanted to create a TIM structure with both chains open and one structure with both chains closed. Therefore, we performed least square fit (LSQ) of backbone of 5TIM chain-A (open) on the chain-B excluding residues K12, H94, E166, loop-6 and loop-7. Same procedure was performed using 6TIM crystal structure where closed chain-B was fitted on open chain-B and excluding the groups mentioned above to create a TIM dimer with both chains closed. The difference between the fitted chains and crystal structures was ~0.03 nm i.e. they very highly identical. These structures would be referred to as fitted-5TIM (F5TIM), fitted-6TIM (F6TIM) and fitted-1R2R (F1R2R). 1NEY is the only crystal structure with the natural substrate DHAP in both the active site pockets. We removed the DHAP molecule from both the active sites to create an apo 1NEY structure.

In all simulations we have chosen cysteines to be neutral (CYN) and lysine’s were protonated (LYP). Histidine’s were either HIE (hydrogen on epsilon nitrogen) or HID (hydrogen on delta nitrogen) depending on the local hydrogen bonding possibilities. The catalytic residue H94 is with hydrogen at epsilon nitrogen^23,43,44^ .

Each crystal structure was placed in a dodecahedron box with a minimum distance of 1nm between the protein and the box walls. The box was solvated using the genbox module and ions were added to neutralize the system. Before the start of the MD simulation, the system of interest is subjected to an energy minimization procedure using steepest descent method until the largest force acting on the system was smaller than 239 kcal.mol^−1^ .nm^−1^ . Energy minimization is performed to remove any unfavorable interactions (like overlapping Van der Waals spheres) present in the crystal structure which may result in large forces and create instability in the system.

After energy minimization, system is slowly equilibrated to the chosen reference temperature and pressure values, which in our simulations are 298 K and 1 atm by coupling the system to a thermostat and a barostat. Temperature and pressure equilibration are done in two phases. First the temperature of the system is brought to 298 K in 100 ps by coupling the system to berendsen thermostat^45^. During the temperature and pressure equilibration solvent is allowed to equilibrate and the protein atoms are position restrained using a force constant of 239 kcal.mol^−1^ .nm^−2^, to preserve the crystal structure.

The end structure of the temperature equilibration is used for pressure equilibration using berendsen barostat for 1 ns. The end structure of pressure equilibration was used for the production run where the protein is free to evolve at constant temperature and pressure of 298 K and 1 atm pressure. Multiple trajectories were started using random starting velocities. Unless mentioned otherwise, same simulation set up and equilibration protocol was used for all the simulations.

- Five trajectories each of F5TIM and F6TIM were performed for 1 μs each.
- Three simulations of apo 1NEY were performed for 300 ns each.
- Three trajectories with F1R2R structure lasting 100 ns each.
- Simulations of unperturbed 5TIM, 6TIM and 1R2R crystal structures were also performed.

### 2.2 Holo TIM simulations

We have performed holo TIM simulations using F5TIM, F6TIM, 6TIM, 1NEY and 1N55^46^ crystal structures. Simulations with DHAP were performed using F6TIM, F5TIM, 6TIM and 1NEY crystal structures. 1N55 is the highest resolution (0.82 Å) structure available for TIM with 2PG bound in the active site pocket. Serena Donnini created the dimer of 1N55 crystal structure by performing a matrix transformation of the crystal coordinates using the BIOMT records provided in 1N55 pdb file. Simulations with inhibitor 2PG and reaction intermediates EDT1, EDT2 were performed with this 1N55 dimer structure. EDT1/EDT2 were placed in the active sites of 1N55 by least square fitting the respective reaction intermediate onto the 2PG present in the crystal structure. For DHAP simulations, the parameterized DHAP was first least square fitted to the substrate analogue G3P present in the chain-B of 6TIM. This structure was used as the starting structure to create DHAP ligated dimers of F5TIM, F6TIM and 6TIM.

After creating the various starting structures with DHAP/EDT1/EDT2/2PG, the same equilibration procedure was applied which was used for apo TIM simulations with one extra equilibration step we call *‘Ligand Equilibration procedure (LEP)’*. In LEP position restraints on both the protein and the ligand were coupled to a lambda (λ) parameter and as λ is transformed from 0 to 1 the force constant of position restraints decreases from 239 kcal.mol^−1^.nm^−2^ to 0 kcal.mol^−1^ .nm^−2^, allowing the protein and ligand to equilibrate. The end structure from this equilibration procedure was used for production run.

### 2.3 Ligand parameterization

TIM interacts with multiple ligand molecules. DHAP and GAP are the natural substrates; 2PG is the strongest binding inhibitor for TIM (**Supplementary Table 1**). Charge on DHAP is −2 and 2PG, EDT1 and EDT2 have a charge of −3^15,22,38,47-49^. EDT1 is the first reaction intermediate when DHAP is converted to GAP, formed after the abstraction of proton from C1 carbon by catalytic base E166. EDT2 is the last reaction intermediate which is awaiting a proton transfer from E166 carboxylate moiety onto the C2 carbon completing the formation of GAP^38^.

As stated earlier General Amber forcefield (GAFF) parameters for these ligand molecules were generated (**Supplementary Figure 2**) using Gaussian03^50^ and Antechamber^51^ package. The structure optimization and charge derivation was done using the basis set mentioned in **Supplementary Table 1**. Antechamber^51^ tool was used to do the RESP^52^ charge fitting for every ligand molecule based on the charges obtained from QM calculation. Generally 6-31g(d)^53,54^ basis set with Hartree-Fock (HF) method is used for charge calculation in GAFF forcefield but this basis set placed negative charges on methyl hydrogen’s in the ligand.Therefore we tried multiple basis sets with HF and B3LYP^55-57^ and B3LYP/6-311G(d)^58,59^ gave us the best charges for DHAP, EDT1 and EDT2. We tried the same for 2PG and finally reasonable charges were obtained with basis sets mentioned in **Supplementary Table 1**.

To test whether the torsional parameters obtained at QM level hold in Molecular Mechanics (MM) forcefield, we performed a potential energy surface (PES) scan around the dihedrals formed by C1-C2 and C2-C3 of bonds of DHAP molecule at QM level using Gaussian03 package and then repeated the same at MM level using Gromacs 4.0.x and if different were optimized to reproduce the QM potential energy surface. Torsional parameters obtained from DHAP for dihedral angles formed by C1-C2, C2-C3 were used for C1-C2, C2-C3 bonds of EDT1 and EDT2 as well. Parameters for DHAP, EDT1 and EDT2 and 2PG are listed in Supplementary information.

The parameterized ligand molecule was placed in the active site of the respective crystal structure and this structure was used for further holo TIM simulations.

- Three trajectories each for F5TIM, F6TIM, 6TIM with DHAP and 1N55 with 2PG, 1N55 with EDT1 and 1N55 with EDT2 in the active site were simulated.
- 1N55, EDT1 and EDT2 simulations were simulated for 300ns each, whereas simulations with DHAP were stopped as soon as DHAP escaped from both the pockets, which was approximately 200ns to 300ns.

### 2.4 Crystal simulations

Surprisingly in our apo TIM simulations with F1R2R and unperturbed 1R2R crystal structures, loop-6 opens up in first 20 ns of simulation time. To further explore the affect of crystal environment on dynamics of loop-6 and loop-7 we performed crystal unit cell simulations as described by Cerutti et al.^60,61^ . One unit cell of 1R2R crystal structure was simulated to see the effect of the crystal environment on dynamics of loop-6 and loop-7. In 1R2R crystal structure each asymmetric unit is made up of two dimers and four asymmetric units form one unit cell. Each unit cell contains 12 open chains and four closed chains. Symmetry records from the pdb file were used to create a unit cell. Such a unit cell contained crystal waters and the protein. Two sets of apo crystal simulations were performed. One structure with 1.1 M DMSO and 0.2 M of MgCl_2_ salt concentration used in mother liquor was created (Crystal Solution Simulation: CSS) and a second simulation with just solvent (Water Crystal Simulation: WCS). Both systems were neutralized by adding additional ions.

A solvent box of the same dimensions as unit cell was created and the unit cell with protein and crystal waters was embedded into the solvent unit cell using g_membed^62^ tool. The resultant structure was subjected to energy minimization as done for apo and crystal simulations, after which temperature equilibration of 1 ns and pressure equilibration of 20 ns was performed. As suggested by Cerutti et al.^60,61^, after pressure equilibration, volume of the simulation box was calculated and if it differed from the crystal unit cell volume by more than 0.3%, solvent molecules were added or deleted from the system.

The whole procedure of energy minimization, temperature equilibration and pressure equilibration were performed until the unit cell with required X, Y and Z dimensions were obtained within ~0.3% of the crystal unit cell volume. The final equilibrated structure contained ~102300 atoms and was used for production run. In all the production run trajectories unit cell volume does not differ by more than 0.3% of the crystal unit cell volume. We have simulated five trajectories with the same equilibrated structure and random starting velocities. All the open chains are combined into Crystal simulations open ensemble (CS-O) and closed chains into crystal simulations closed ensemble (CS-C).

We have also performed holo crystal simulations with 1N55 crystal structure with 2PG bound to the active site pocket. Starting structure for 2PG crystal simulations (CS-2PG) was subjected to the same protocol as the 1R2R crystal simulations. In 1N55 crystal structure, there is one monomer per asymmetric unit i.e. four monomers per unit cell and we have simulated one unit cell in MD. The 1N55 unit cell structure was subjected to similar equilibration procedure as 1R2R unit cell structure to create the simulation box which differed in volume by no more than ~0.3% of X-ray crystal unit cell. The end structure of volume equilibration was used to start three production run simulations. The volume equilibrated was also subjected to LEP and this structure was used for three more production run simulations.

- Five apo TIM crystal unit cell simulations of 300 ns length, with the corresponding equilibrated structures of WCS and CSS (which were created from 1R2R crystal structure) were performed.
- Six simulations with 2PG ligated 1N55 crystal structure (CS-2PG) three with the volume equilibrated structure and three with the end structure of LEP were performed.
- The reference temperature for unit cell simulations of 1R2R and 1N55 was 291 K^42^ and 295 K^46^, the room temperature at which these proteins were crystallized.

TIP3P water model was used for solvent^63^. We have used Particle-mesh Ewald algorithm^64,65^ to evaluate the electrostatics interaction with a cutoff of 1.0 nm. Van der Waals interactions were calculated with a cut-off of 1.4 nm. LINCs^66^ algorithm was used to constrain on all-bonds. Nose-Hoover (NH) thermostat^67,68^ and Parrinello-Rahman (PR) barostat^69,70^ were used for maintaining NPT ensemble for the production run simulations. For unit cell simulations we used Berendsen barostat^71^ along with because if its efficiency in maintaining unit cell volume better than PR barostat along with v-rescale thermostat^72^. Coordinates were saved every 20 ps and energies were written every 10 ps.

## 3 Results

### 3.1 Analysis of various X-ray structures

From the 126 TIM X-ray structures available, 93 were analyzed using **X-ray P**DB **A**nalysis **T**ool (XPATv1.2) written by us which reads a pdb or multiple pdb files and calculates distances or angles or dihedral angles for selected residue(s), as requested by the user. Twenty-three structures with resolution greater than 2.5 Å were excluded from the analysis. Phi and psi angles of loop-7 residues Y209 to N214 and loop-6 residues E166 to T178 were calculated to identify the dihedral angles undergoing a change between open and closed states (**Supplementary Table 2**).

As discussed in section 1.2, dynamics of V168 from loop-6 N-terminus and C-terminus residues K175 and T178 were monitored in NMR experiments^15,22^. So, to segregate TIM X-ray structures into open and closed states based on N-terminus and C-terminus conformations of loop-6, distance between W169 from N-terminus of loop-6 and loop-7 residue S212 was calculated in all X-ray structures (W169-S212). And to determine loop-6 C-terminus conformation, distance between loop-6 C-terminus residue V176 and S212 (V176-S212) was calculated. Based on W169-S212 and V176-S212 distances (**Supplementary Table 2**), TIM structures were classified as open TIM (OTIM) when both loop-6 N and C-terminus are open (W169-S212: 1.15 nm, V176-S212: 0.88 nm) and as closed TIM (CTIM) when both N and C-terminus are closed (W169-S212: 0.95 nm, V176-S212: 0.50 nm).

As discussed in section 1.3, loop-7 also exists in open and closed conformations. Loop-7 is segregated into open and closed conformations based on the backbone phi (Φ) and psi (Ψ) dihedral angles of residues G210, G211, S212 and V212. Dihedral angles G210Φ, G210Ψ, G211Φ, G211Ψ, S212Φ, S212Ψ and V213Φ differ by more than 25° between open and closed conformations (**Supplementary Table 2**).

Majority of X-ray structures (**Supplementary Table 3**) in the absence of ligand are in open state (43) and in the presence of ligand are in closed state (73). It must be noted that there are two crystal structures with loop-6 in closed conformation (1R2R-B, 2VEN-A^73^) without the ligand in the active site pocket. Loop-7 of 1R2R-B is also in closed conformation whereas in 2VEN-A, excluding S212Φ, entire loop-7 is in open conformation.

In 1R2R crystal structure, out of four chains A, B, C and D, chain-B is in closed state. There is no ligand in the active site but there is a water molecule (W171) which is present at the location where phosphate moiety of ligand is situated forming a hydrogen bond with S212 amide group^42^. Further studies are necessary to ascertain, if a water molecule located at the phosphate moiety’s position can cause flip of loop-7 to closed state. Closed loop-6 of 1R2R-B was reported as a feature of TIM’s conformational heterogeneity by Ricardo et.al^42^ .

The second crystal structure which has loop-6 in closed conformation is 2VEN. Loop-6 in chain-A of 2VEN dimer adopts closed conformation in the absence of ligand. In chain-A of 2VEN-A electron density for lid region (I172-GT-175G) of loop-6 is missing. In OTIM loop-6 has high B-factors and same was observed for loop-6 in 2VEN-A^73^. But based on loop-6 N and C-terminus distance we would suggest that 2VEN-A as a closed structure. These two closed-apo structures suggest that loop-6 might also exist in closed conformation even in the absence of the ligand, though open state is the preferred conformation in apo TIM.

In our analysis we also came across 14 chains where both loop-6 and loop-7 were in open conformation with ligand occupying the active site (open-holo). To understand the reason behind these structures being in open conformation despite of the occupied active site, we analyzed the ligand orientation. Each structure was visually compared with the encounter complex (EC) structure of 1LZO-B^74^ and reactive complex (RC) of 1NEY^75^, 1N55^46^ structures (**Supplementary Figure 3**). In 1M7O^10^ (3PG: 3-Phosphoglycerate) and 1M7P^10^ (G3P: Glycerol-3-Phosphate) where substrate analogs are bound in RC state, the S96F mutation prevents loop-6 from closure. Rest of the 10 chains had ligand in the encounter complex position. When we analyzed the orientation of ligand in the closed-holo structures we found out that all the closed-holo chains had the ligand in RC orientation.

In the encounter complex we found van der Waal’s clashes between the phosphate moiety of the inhibitor and the loop-6 residues I171 and G172 which can prevent loop-6 closure. The mechanism of loop-7 conformational change and influence of ligand on loop-7 has not been explored which we address in this work in section 3.5.2.1. As stated earlier, in chain-B of 1R2R crystal structure, a water molecule occupies the space of one of the oxygen’s of the phosphate moiety, within hydrogen bonding distance G172 of loop-6 and S212 of loop-7. Looking at the X-ray structures it might be speculated that the electrostatic repulsion between the phosphate moiety and the carbonyl oxygen might drive the flip of G211Ψ and S212Φ to closed conformation and it is quite surprising that a 0.2 nm change in shift of phosphate moiety towards the mouth of active site induces a loop-7 opening.

Our analysis also revealed structures where loop-6 N and C-terminus were in different states i.e. both the hinges were neither closed nor open at the same time (**Supplementary Table 4**). Crystal structures 2BTM , 2J24 ^76^ are mutant TIM structures whereas 2VEI^73^ is a monomeric TIM variant. The P167A mutation in 2J24 induces this conformational heterogeneity in loop-6 whereas in 2BTM the mutations H12N, K13G are not in loop-6. In 2VEI, V233 was mutated to A233 to extend the active site and enable binding of citrate (CIT) molecule. Loop-6 and loop-7 are in closed state, in these CIT bound structures. The mutant structures have decreased catalytic activity^76,77^. 1WYI is a wild type human TIM with no mutations at 2.20Å resolution^78^.

To summarize the TIM X-ray structures analysis:

- In crystal structures loop-6 and loop-7 exist in open conformation in the absence of ligand.
- Loop-6 may also exist in closed conformation in the absence of the ligand (2VEN-A and 1R2R-B).
- Apart from chain-B of 1R2R X-ray structure where a water molecule is located in the place of phosphate moiety, loop-7 does not exist in closed conformation when there is no ligand bound to the active site.
- There is only one crystal structure with natural substrate DHAP bound in the RC orientation with both loop-6 and loop-7 in closed conformation.
- Ligand molecules i.e. substrate or inhibitor can exist in RC and EC states in the active site.
- Loop-6 and loop-7 are in open conformation when the ligand is in the EC state.
- Only when a ligand molecule is in the RC state, both loop-6 and loop-7 are seen in closed conformation.

### 3.2 Root mean square deviation

We have looked at the Root mean square deviation (RMSD) values of all the trajectories. RMSD tells us how much the structures deviate from the starting crystal structure. All the trajectories showed a stable RMSD of less than 0.2 nm. Major structural changes are seen in loop-6 with minor fluctuations in loop-5 and loop-7. In rest of the protein there are no significant structural changes.

### 3.3 Ligand residence time

We have simulated OTIM and CTIM structures with natural substrate DHAP, reaction intermediates EDT1 and EDT2 and inhibitor 2PG. In each holo TIM simulation we have calculated the time spent by each ligand in RC, EC and apo orientations which we define as residence time (RT). RT is calculated using Principal component analysis (PCA) and RMSD analysis. To describe the change in orientation of each ligand in the active site pocket from RC to EC orientation, we have used the eigenvector-1 (LEv-1) from PCA. PCA was performed on chain-B of 1LZO where 2PG is sitting at the mouth of the active site which is EC orientation (**Supplementary Figure 3a**) and 1NEY structure where DHAP is present in RC orientation. LEv-1 describes the ligand transition from RC to EC state.

We also calculated RMSD of the ligand molecule (excluding hydrogen atoms) with respect to the ligand molecule in the starting structure of the simulation by using the protein backbone as the fitting group. When the ligand moves out of EC state into the solution, RMSD of the ligand with respect to the ligand molecule in the starting structure is greater than 0.7 nm. Therefore, all the frames of the trajectory where RMSD of the ligand is less than or equal to 0.7 nm, are used to calculate RT. Frames with RMSD greater than 0.7 nm are considered apo.

Based on eigenvalues of LEv-1 we classify ligands orientation into RC (−0.50 nm to 0.50 nm) and EC (−0.5 nm to 1.0 nm) states. We have started all our simulations with DHAP, 2PG, EDT1 and EDT2 in the RC orientation. In **Figure 6** we have plotted the average RT of each ligand molecule over the entire ensemble and the error bars were calculated by bootstrap method which is described in supplementary information.

**Figure 6:**
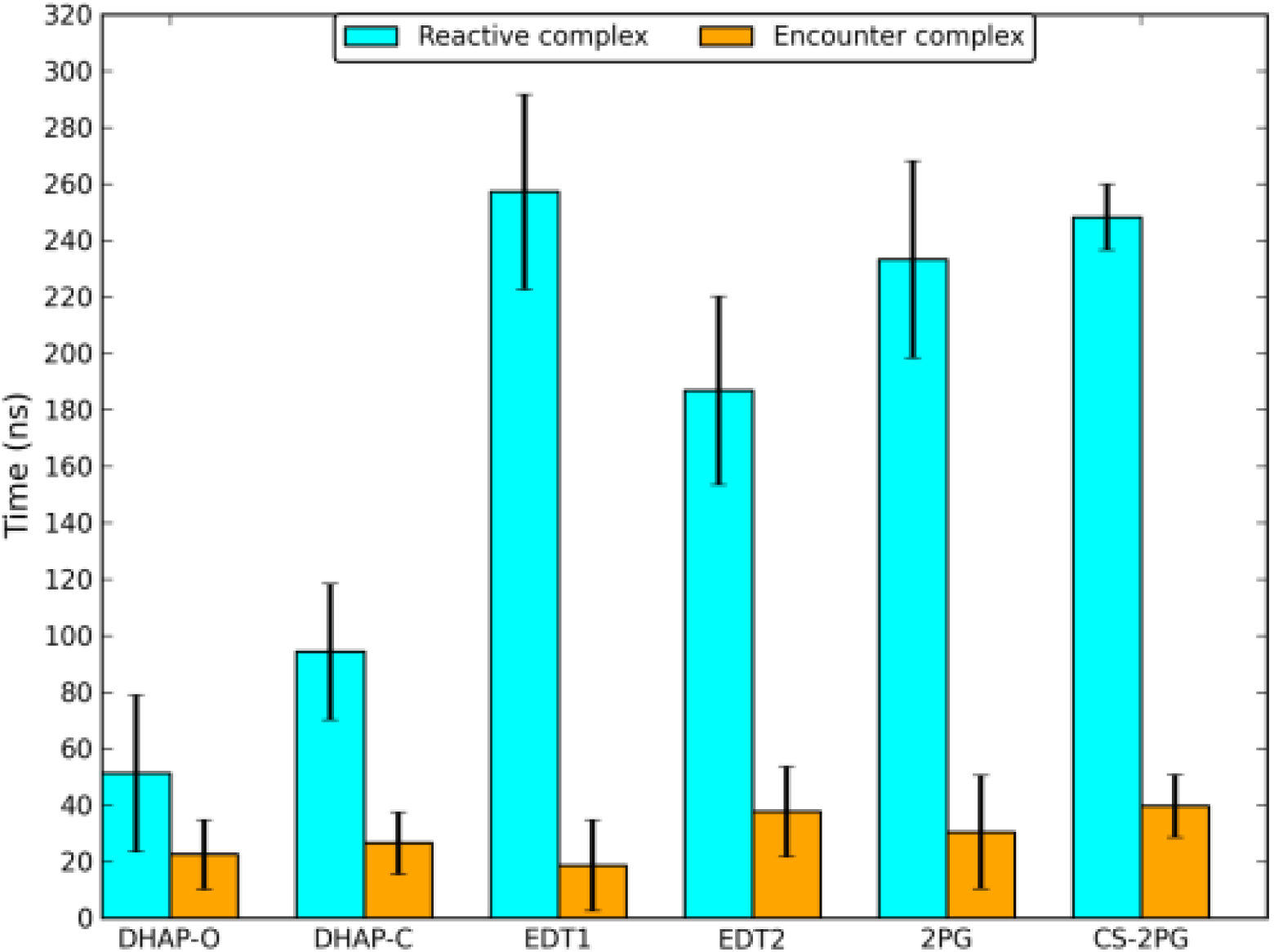
The average residence time (RT) of natural substrate DHAP, reaction intermediates EDT1, EDT2 and inhibitor 2PG in normal MD simulation (2PG) and crystal simulation (CS-2PG) the active site pocket of TIM, in reactive complex (RC) and encounter complex (EC) orientation. Error bars were calculated by bootstrapping the averages.

As seen in **Figure 6** all the ligand molecules spend more simulation time in RC state compared to the EC state. We have placed DHAP in both OTIM and CTIM active site pockets. Irrespective of the starting TIM conformation, DHAP moves out of the pocket in all the simulations. The average RT (time spent in RC+EC orientations) of DHAP with OTIM as the starting structure is 74 ns which is less compared to 121 ns when the simulation starts with CTIM (**Figure 6** ).

Inhibitor 2PG, in five out of six chains is bound to the active site pocket for entire 300 ns of simulation. Only in one chain out of the six chains we observe 2PG moving into the solution after 117 ns (data not shown). Contrary to X-ray data, in holo TIM simulations with DHAP and 2PG loop-6 does not stay closed and transiently samples open and closed conformations (which is discussed further in 3.4.1). So we have also performed simulations with the reaction intermediates EDT1 and EDT2 (**Supplementary Figure 2**).

Similar to 2PG, EDT1 stays inside the active site pocket for entire length of simulation for 300 ns in five out of six chains and only in one chain EDT1 moves into the solution after 157 ns. Whereas in the simulations with EDT2 bound to the active site, we observe EDT2 dissociating into the solution in all our simulations. Average RT of EDT2 (224 ns) is much longer than the natural substrate DHAP and shorter than EDT1 (275 ns; **Figure 6** ). In comparison, 2PG, EDT1 and EDT2 spend more time in the active site compared to the natural substrate DHAP (**Figure 6** ).

TIM has achieved maximum theoretical efficiency possible for an enzyme (10^9^ M^−1^ .s ^−1^) and is limited by the diffusion of the substrate into the active site^1^. To achieve this turnover rate not only would the enzyme optimize the efficiency of catalytic steps but would also get rid of the product formed in the active site as soon as possible. Taking that view point into consideration it would be of no surprise that DHAP escapes into solution in all our simulations.

2PG is a transition state analogue and the charge of −3 compared to the charge on substrates DHAP and GAP of −2, provides additional electrostatic stabilization in the active site contributing to its higher binding affinity^79^. Hence 2PG should bind very strongly to the enzyme and 2PG stays bound to the active site for the entire 300 ns of the simulation. Enzymes are speculated to stabilize the reaction intermediates more compared to the substrate, and we observe higher RT for reaction intermediates in our simulations compared to the natural substrate. But to confirm our speculations we need to calculate the binding free energy of these ligand molecules in OTIM and CTIM conformations which is a more reliable measure compared to the RT presented here, will be the focus of our future work.

### 3.4 Dynamics of loop-6

Loop-6, as discussed in section 1.2 plays a very pivotal role in TIM’s function. The fundamental questions we want to answer about loop-6 are as follows.

- Nature of loop-6 conformational change, is it ligand driven or a spontaneous change driven by thermal fluctuations?
- Is loop-6 conformational change a rigid body motion i.e. both N and C-terminus move together (correlated) or N and C-terminus of loop-6 independent?
- What is the free energy difference between the open and closed loop-6 in both presence and absence of the ligand, and what factors contribute to this free energy difference?
- How does binding of a ligand (substrate/inhibitor/reaction intermediates) affect dynamics of loop-6?

To understand the various aspects of dynamics of loop-6, we performed multiple apo and holo TIM simulations. In our holo TIM simulations we have used the natural substrate DHAP, inhibitor 2PG and reaction intermediates EDT1 and EDT2. All the ligated TIM simulations start with ligand in the RC orientation.

#### 3.4.1 Apo and holo TIM simulations

We have performed 16 apo TIM simulations in which there are 13 OTIM chains and 19 CTIM chains. Out of 13 OTIM chains with open loop-6 conformation, in 10 chains, 2 to 14 transitions between open and closed conformations of loop-6 are observed. In three OTIM chains there are no transitions. All the simulations that start with loop-6 in closed conformation, undergo a conformational change to open state in the first 20 ns of simulation time (**Figure 7**), longest being 150 ns. In seven CTIM chain trajectories no transitions between open and closed loop-6 are observed and 2 to 21 transitions are seen in rest of the 12 CTIM chains. A typical loop-6 N-terminus (W169-S212) and C-terminus (V176-S212) distance plot is shown in **Figure 7** where we have 20 transitions between open and closed conformations starting with a CTIM structure.

**Figure7:**
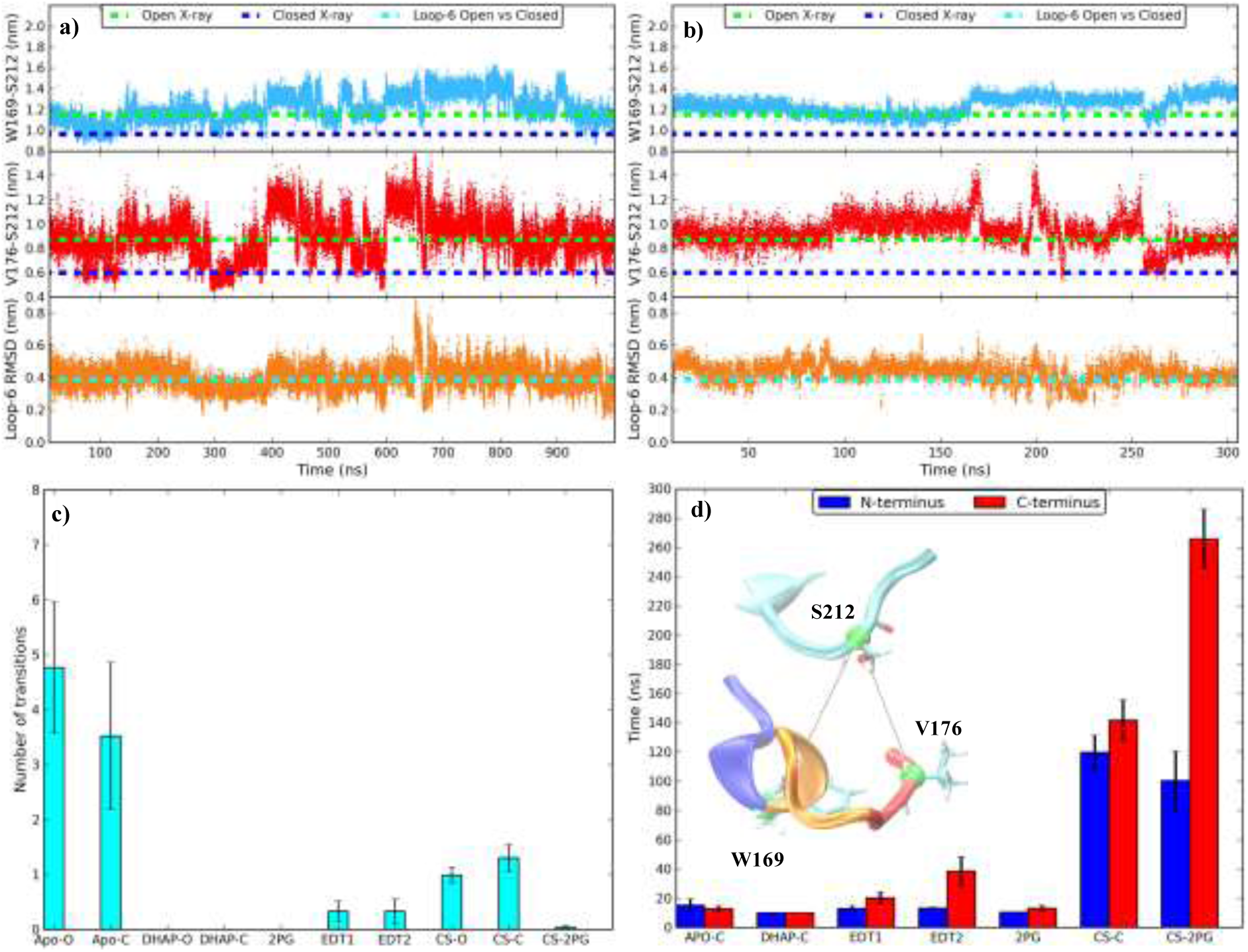
Dynamics of Loop-6 N-terminus (W169-S212) and C-terminus (V176-S212) in a microsecond trajectory which started with an apo CTIM structure (a) and 300 ns long trajectory where EDT1 is bound for entire length of simulations (b). Loop-6 RMSD with respect to closed loop-6 is shown in the third panel. c) Average number of transitions between open and closed conformations of loop-6. For ligated ensembles only that part of the trajectory during which ligand is bound to the active is considered. d) Average time taken by loop-6 N-terminus (W169-S212) and C-terminus (V176-S212) to undergo conformational change from closed to open conformation for the first time in the simulation when the simulation started with a CTIM structure, in various ensembles. Error bars are calculated using bootstrap method. In inset of plot (d) loop-6 with N-terminus (blue), tip (orange) and C-terminus (red), loop-7 (cyan) are shown.

To check if the closed loop-6 in MD simulations resembles the closed loop-6 of the crystal structure, we have calculated the RMSD of the backbone of loop-6 with respect to loop-6 of the CTIM X-ray structure for all our trajectories. The entire protein backbone excluding loop-6 is used as the fitting group. The RMSD difference observed between MD loop-6 and X-ray structure is between 0.15 nm and 0.2 nm (**Figure 7a** **&** **7b**).

Our analysis of TIM X-ray crystal structures has revealed that when ligand is bound in RC orientation loop-6 is in closed conformation. Surprisingly ,in our holo TIM simulations with DHAP/2PG/EDT1/EDT2 in RC/EC orientation no preference for closed conformation of loop-6 is observed (**Figure 7b**) as expected from the X-ray data. In all the holo TIM simulations that start with CTIM structure, loop-6 undergoes a conformational change to open loop-6 in the first 40ns (**Figure 7d**). In simulations with DHAP/2PG during the simulation when the ligand is bound to the active site in RC orientation, no transitions back to the closed conformation of loop-6 are observed (**Figure 7c**). In simulations with EDT1 and EDT2 in RC orientation we observe one transition each, in two out of five chains in which ligand is bound to the active site (**Figure 7c**).

Loop-6 to and fro samples open and closed conformations in apo TIM simulations (**Figure 8**). There are fewer transitions in holo TIM simulations because of shorter simulation times (**Figure 8**). It might be possible that the time required for closure of loop-6 is a much slower process than 300 ns of simulation time during which the ligand molecules are bound to the active site, to induce closure of loop-6. As expected from X-ray structure data in holo TIM simulations we do not see preference for closed state. Richard and coworkers^80^ have proposed that the binding energy of the phosphate moiety is used for inducing closed conformation for loop-6 which is not observed in our holo TIM simulations. Similar properties of loop-6 dynamics were observed in the molecular dynamics simulations of yeast TIM by Liao et al.^81^.

**Figure 8:**
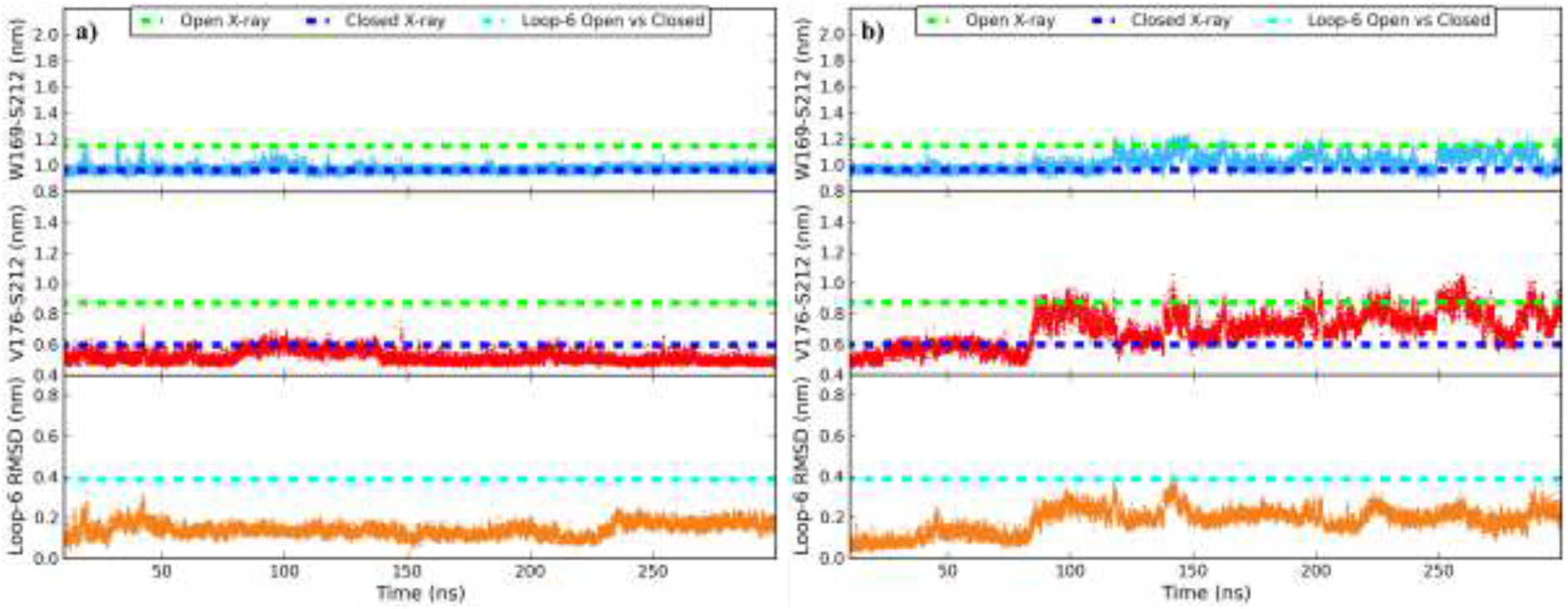
Dynamics of loop-6 in crystal simulations. (a) Simulation which started with closed loop-6 stays closed for the entire length of simulation and (b) simulation with four transitions between open and closed conformations of loop-6. Loop-6 RMSD with respect to closed loop-6 is shown in the third panel.

#### 3.4.2 Effect of crystal environment

In apo and holo TIM ensembles when simulations start with CTIM structure loop-6 undergoes a conformational change to open state in the first 40 ns of simulation (**Figure 7d**). To see if there was an influence of crystal environment on the conformation of loop-6 i.e. may be only in the X-ray crystal, closed conformation of loop-6 is stable. So, we performed both apo and holo crystal unit cell simulations with 1R2R (apo) and 1N55 (holo with 2PG in active site pocket) structures. There is no difference observed in the behavior of loop-6 in the crystal solution simulation (CSS) and water crystal simulation (WCS) hence we discuss both of them as one ensemble. In crystal simulations CS-C and CS-2PG, time taken for this conformational change is greater than 100 ns (**Figure 7d**) compared to the first 40 ns in normal MD simulations. Time taken by N-terminus and C-terminus of loop-6 to open is different with N-terminus opening first followed by C-terminus. The reason behind this is addressed in section 3.4.4.

Diffusion constant of water from MD simulations was calculated using Einstein relation^82^ from mean square displacement of water molecules. Solvent dynamics in crystal simulations is 10 times slower than in the normal MD simulations (**Supplementary Table 5**). Because of the slow solvent dynamics, the dynamics in the crystal simulations is slow (**Figure 8**). In 9 out of 40 chains, loop-6 stays closed for the entire 300 ns of simulation (**Figure 8a**). In six chains, C-terminus of loop-6 stays closed and N-terminus undergoes transitions between open, closed structures and the opposite is seen in only two chains. Even in the 2PG ligated unit cell simulations though we observe a slower dynamics of loop-6, i.e. loop-6 undergoes conformational change from closed state to open state. It is clear that the crystal environment slows down dynamics of loop-6, nonetheless loop-6 does undergo a gradual conformational change in both apo and holo crystal simulations (**Figure 8b**).

Diffraction data for 1R2R and 1N55 was collected at 100 K. It is a common practice to flash freeze the protein crystal to less than 125 K to reduce the radiation damage to the crystal^83^. An assumption made prior to flash freezing a protein is that this procedure traps the protein in conformation at the room temperature and does not affect the conformational equilibrium of the protein^84-86^ .

Time scale of this procedure is in seconds and there is uncertainty about the effect of flash freezing on conformation of the crystallized protein^87-90^. Halle^87^ suggested that during flash freezing procedure, different regions of protein may get trapped in low a enthalpic minimum at different temperatures and the whole structure may not be a representative of structure at one single temperature. Dunlop et al.^88^ in 2005 collected diffraction data for six crystals, three at 100 K and three at 293 K and demonstrated that there is a significant difference with respect to side chains and hydrations sites specifically at the surface of the protein.

Our crystal simulations were performed at 291 K. Simulations at 100 K will further slow down the dynamics and if the assumption that conformational equilibrium is not altered by flash freezing, our simulations at 291 K should reproduce the experimental data. To verify this, we have compared the B-factors of C-alpha atoms from crystal open (CS-O) and crystal closed (CS-C) MD simulations with experimental B-factors of chain-A from 1R2R (OTIM Crystal @ 100 K), chain-B from 1R2R (CTIM Crystal @ 100 K), 1R2S (OTIM Crystal @ 298 K) and 1N55 crystal structures for which diffraction data was collected at 100 K, 298 K and 100 K respectively (**Supplementary Figure 4a**).

We calculated the spearmen rank correlation coefficient between the B-factors from MD crystal simulations and B-factors from crystal structures and B-factors calculated from normal APO MD simulations. Crystal MD simulations were able to reproduce the trend seen in X-ray crystal structures and show a moderately strong correlation between 0.60 and 0.78 (**Supplementary Table 6**). Residues 1-10, 38-50, 75-90 reproduce the trend seen in crystal data but have much smaller fluctuations. This may be because they are trapped in some local minimum and require more simulation time to equilibrate.

Protein residues 29 to 32 in CS-O, residues 132 to 142, loop-7 residues G207 and G208 in both CS-O and CS-C have higher B-factors than 1R2R crystal at 100 K but similar to seen at 1R2S. Loop-6 residues 167 to 174 have higher flexibility when compared to crystal data both at 100 K and 291 K. But majority of the protein shows B-factors resembling the 1R2R crystal at 100 K hence higher correlation of greater than 0.7 with 1R2R crystal structure than 1R2S. In both the crystal simulations CS-C, CS-O and normal apo MD simulations fluctuations are seen in the same protein fragments with one exception of residues 69 to 72 (solvent exposed loop) which show higher fluctuations in normal MD simulations (**Supplementary Figure 4a**).

B-factors calculated from 1N55 crystal unit cell simulations (**Supplementary Figure 4b**) also show higher B-factors than the crystal data especially in the loop-6 residues 167 to 178 and loop-7 residues 211 to 217. Rest of the residues which show deviation from crystal data are present in the dimer interface (68 to 75 and 99 to 104) or are present in solvent exposed loops (29 to 34, 52 to 57 and 149 to 156). As each asymetric unit of the 1N55 unit cell has a monomer higher fluctuation in the dimer interface is expected where there are contacts with the other monomer. Though the temperature in 1R2R and 1N55 unit cell simulations was maintained at 288.3±0.01 K and 292.2±0.01 K the agreement with experiment is not exact. Especially we observe deviations in loop-5, loop-6 and loop-7 regions. Loop-5 and loop-7 resemble the crystal data at 298 K.

Aparicio et al.^42^ argued that the closed chain in 1R2R crystal structure is represents the “conformational heterogenity” of the protein which means that loop-6 can adapat open and closed conformations in apo TIM as well. Our simulations support this argument as we see transitions between open and closed conformation of loop-6 in both 1R2R and 1N55 crystal unit cell simulations. The time taken for loop-6 to open starting with a closed conformation is much longer in the crystal enviroment because of the slow solvent dynamics. Authors have ruled out effect of crystal contacts and DMSO on the closed conformations of loop-6 and loop-7 and pointed out to a water (Water 171) which interacts with amide groups of G172 of loop-6 and S212 of loop-7, stabilizing the respective closed conformations.

We have performed simulations at 291 K and 295 K respectively for 1R2R and 1N55 crystal simulations and Aparicio et al.^42^ also suggested that effects of crystallization process cannot be overlooked. To verify this hypothesis that flash freezing affects on the conformation of loop’s 5, 6 and 7, multiple annealing simulations of structures from MD simulations at 298 K to 100 K (at which diffreaction data was collection) in microsecond to millisecond time scale starting with open loop-6 and loop-7 have to be performed which is computationally very intensive.

#### 3.4.3 Loop-6 N and C-terminus correlation

Berlow et al.^22^ and Wang et al.^15^ observed different population ratios for N-terminus and C-terminus hinges of loop-6 and suggested requirement of further studies to explore this novel view of loop-6 dynamics. To verify this, we divided loop-6 into five conformations based on W169-S212 and V176-S212 distances for loop-6 (**Supplementary Table 7**). In all apo and holo TIM ensembles population percentage of each conformation of loop-6 in each simulation was calculated.

If loop-6 N and C-terminus hinges are correlated and loop-6 moves as a rigid body we would expect only O, IMD and C states to be populated. And in all our simulations both apo, holo and unit cell simulations, not only do we observe O, C and IMD states but also different populations for NC and CC states. In **Figure 9**, loop-6 average population percentage in each conformation of loop-6 is shown for four ensembles. The uncertainties in the population percentages were calculated by boot strap method as described in supplementary information.

**Figure 9:**
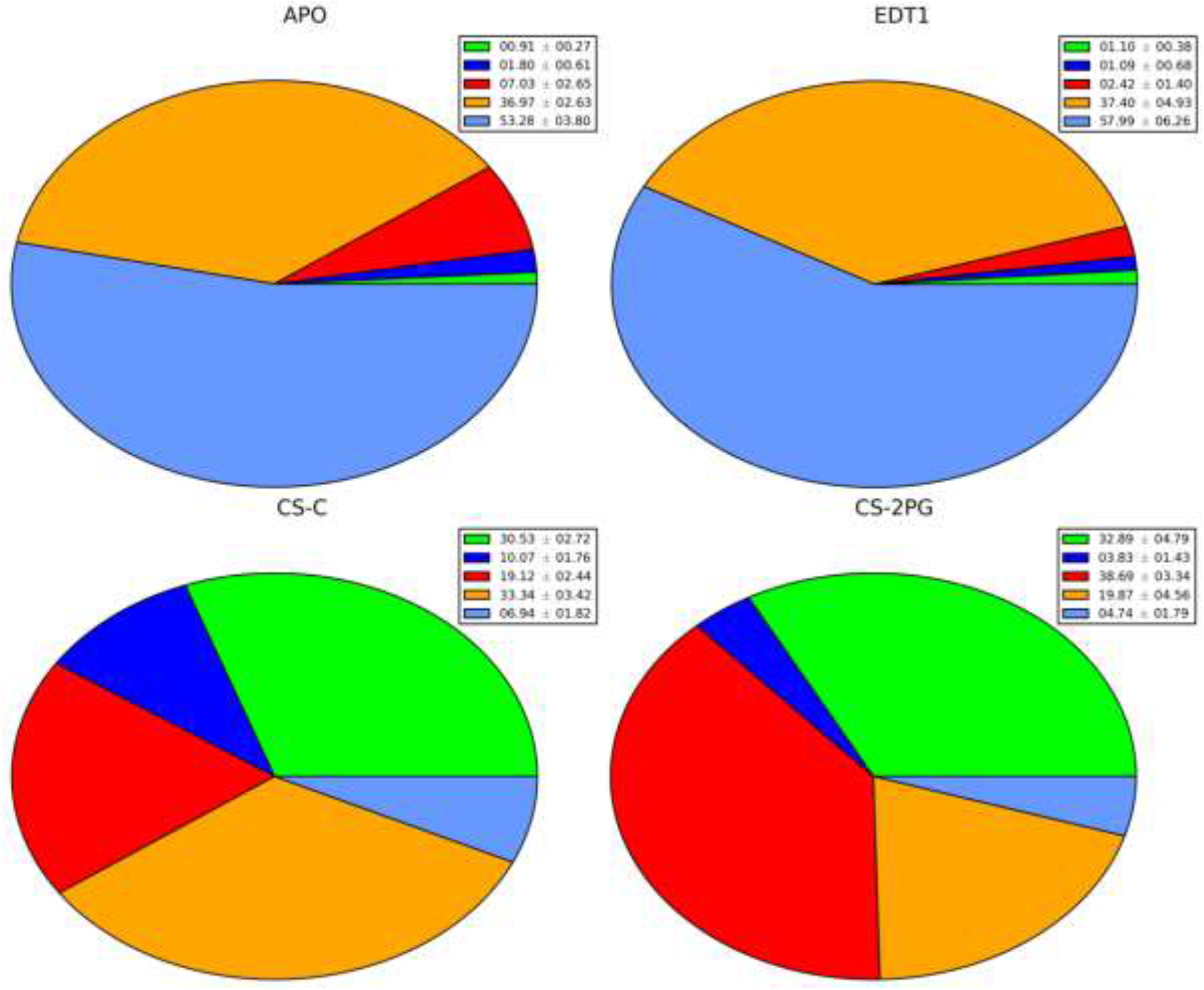
Average population percentages of loop-6 in various conformations, in Apo TIM ensemble (APO), apo unit cell simulation started with closed structure (CS-C), CTIM simulation with EDT1 bound in the active site (EDT-1) and 1N55 unit cell simulation ligated with 2PG. The value in square brackets is the percentage of simulation time 2PG or EDT1 is bound in RC orientation. Loop-6 conformations Open (cyan), closed (green), N-terminus is closed (dark blue), C-terminus is closed (red) and neither N-terminus nor C-terminus are open or closed (orange). The average population percentage with error bar is given in the legend for each pie chart.

As stated earlier in sections 3.4.1 and 3.4.2 we do not see any influence of the ligand bound in the active site on the population of loop-6 in a particular conformation. In apo and holo TIM simulations percentage of closed loop-6 conformation is less than 1% and only in CS-C and CS-2PG it is 30 % and 33% (**Figure 9**). The percentage of NC and CC conformation varies between 1% to 10% and the difference can be attributed to the difference in the apo ensemble (23 μs) and holo TIM ensemble (1.7 μs to 2.7 μs) sizes.

In the NMR experiments loop-6 was only identified as open (NMR-Open: NMR-O) and closed (NMR-Closed: NMR-C) states. Based on the angular hop of W169 present in the N-terminus of loop-6, which was used as an indicator of loop-6 conformation by McDermott and coworkers^18-21^ we have segregated loop-6 five conformations into NMR-O and NMR-C.

We also calculated the normalized mutual information^91,92^ (NMI) between W169-S212 and V176-S212 distances in all the TIM ensembles (**Figure 10**). On a NMI scale of 0 to 1, 0 means no correlation and 1 means perfect correlation. In all our simulations we observe a consistent NMI of less than 0.2 (**Figure 10**), which would imply that N-terminus and C-terminus of loop-6 are uncorrelated. The reason behind this independent movement is that Nterminus of loop-6 interacts with loop-5 and is obstructed by loop-5 before it can open which is discussed in sections 3.4.4 and 0. Whereas C-terminus is solvent exposed and is not inhibited by part of the protein in regards to its dynamics.

**Figure 10:**
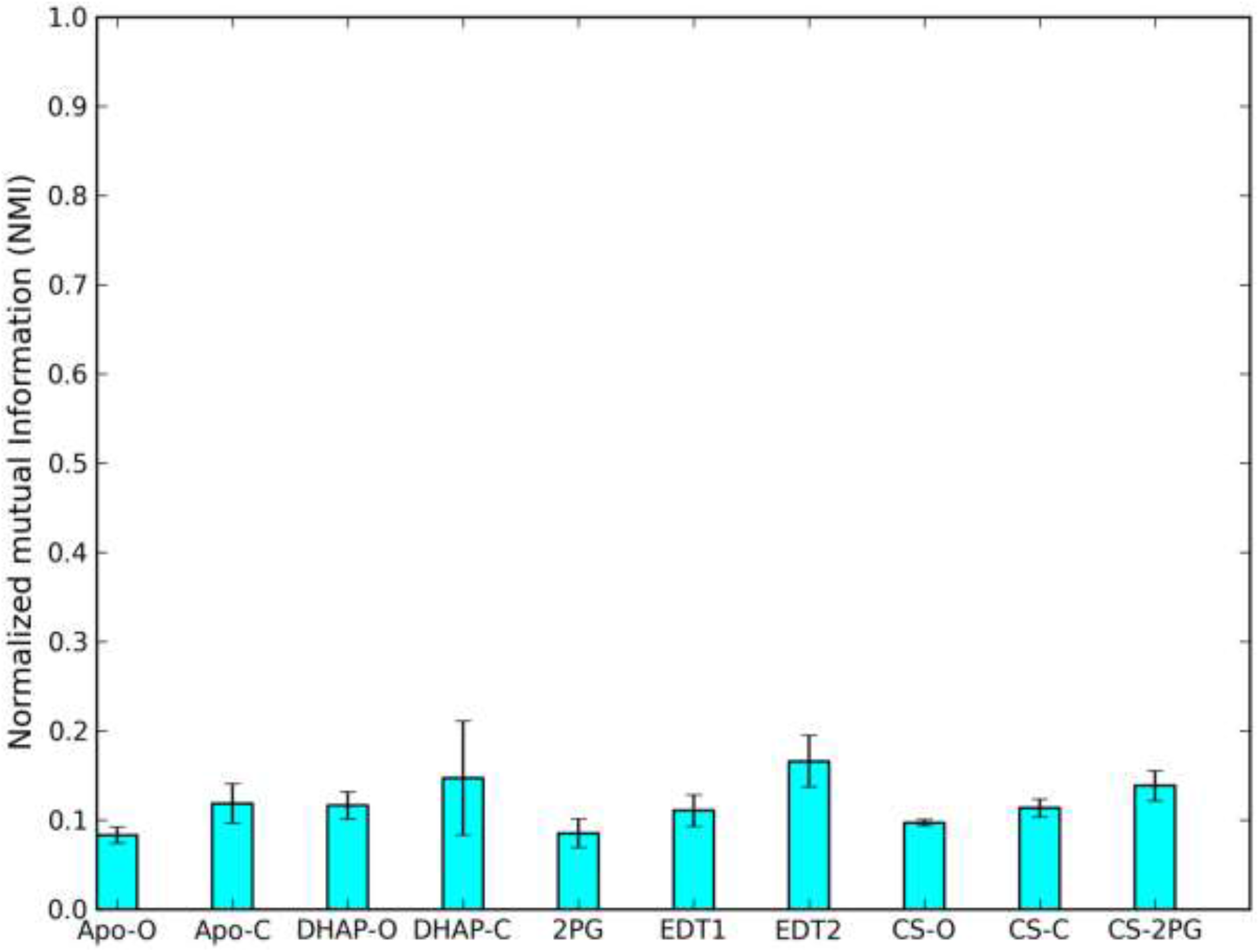
Normalized Mutual Information (NMI) between W168-S212 and V176-S212 distances in various TIM ensembles. On a scale of 0 to 1, 0 means no correlation and 1 means perfect correlation.

#### 3.4.4 Effect of Loop-5 on loop-6 conformation

The effect of loop-5 on dynamics of loop-6 is largely unexplored. A non-zero exchange contribution was detected for loop-5 residues G127 and L130 by Berlow et al. suggesting a conformational change for these residues but this was not further investigated^22^. Only a minor structural change is observed in loop-5 between the OTIM and the CTIM X-ray structures with RMSD less than 0.05 nm. Loop-5 residue E128 forms a hydrogen bond with W169 of loop-6 in closed conformation and this hydrogen bond is broken in open loop-6.

To quantify the correlation existing between loop-5 and loop-6 residues in all the ensembles, we used the generalized correlation coefficient (rMI) developed by Lange et al.^93^ which calculates correlations between atoms based on the atomic fluctuations. Two pairs of atoms have correlated motion will show higher correlation (on a scale of 0 to 1, 0 means no correlation and 1 means correlation) than the atoms whose motion is not coupled. There is a much higher correlation (rMI between 0.48 and 0.60) between loop-5 residues 127 to 131 and loop-6 N-terminus residues P167, V168 and W169 (**Figure 11**). The correlation between loop-5 residues 127 to 131 and loop-6 residues decreases to 0.16 as we proceed towards loop-6 C-terminus residues (**Figure 11**).

**Figure 11:**
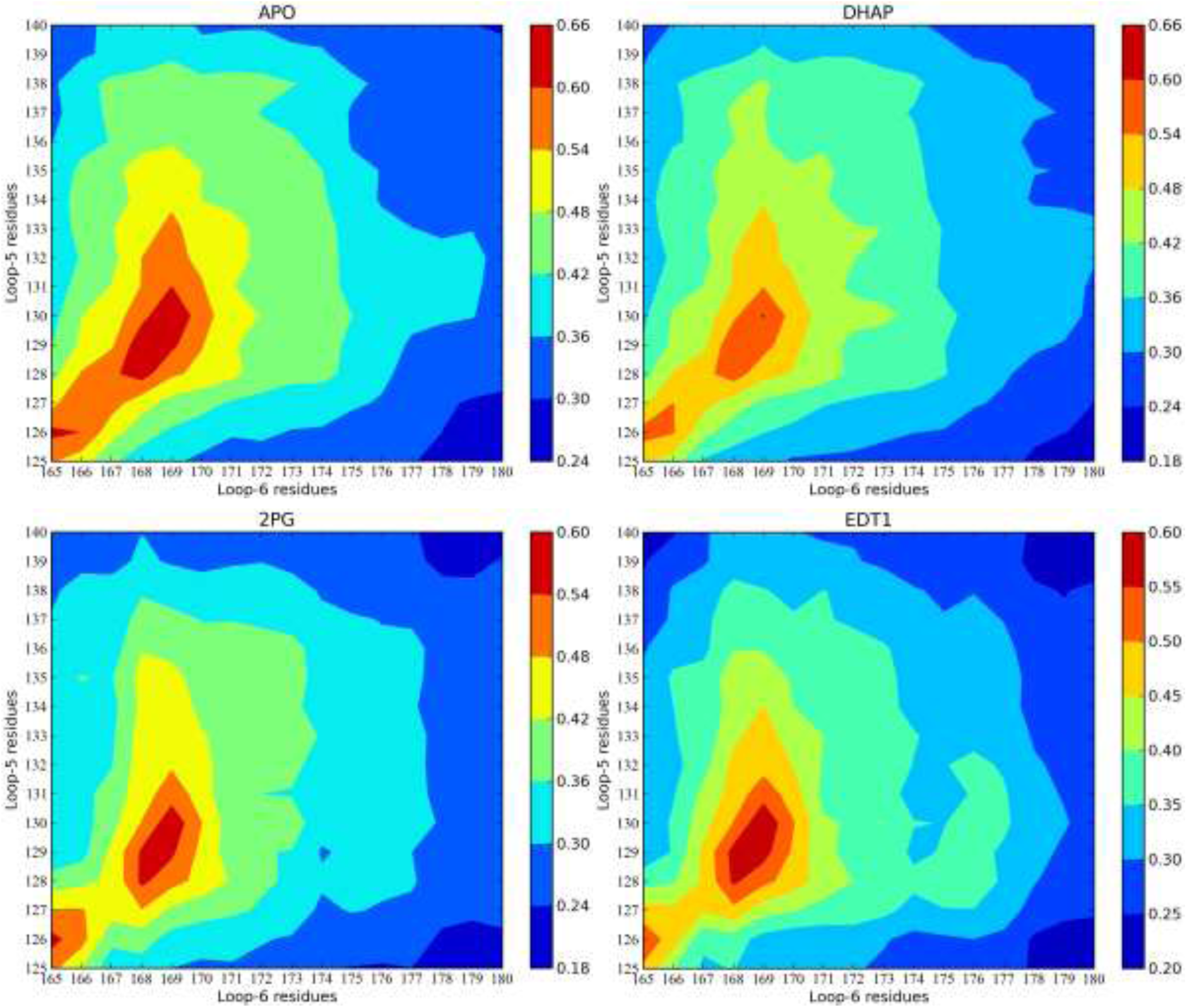
Correlation between Loop-5 residues 127 to 137 and loop-6 residues 167 to 178 in apo TIM ensemble (APO), DHAP ligated TIM ensemble (DHAP), 2PG ligated TIM (ensemble) and EDT1 ligated TIM ensemble (EDT1). Loop-6 N-terminus residues are P167, V168 and W169. C-Loop-6 C-terminus residues are K175, V176 and A177. On a scale of 0 to 1, 0 means no correlation and 1 means perfect correlation.

The correlation between loop-5 residues 127 to 131 and loop-6 C-terminus residues 175 to 177 is less than 0.35. And the nature of correlation does not change in presence of DHAP or EDT1 or 2PG or in unit cell simulations (**Figure 11**). Our results show that loop-6 Nterminus hinge is more correlated with loop-5 residues 127 to 131 then loop-6 C-terminus hinge.

We performed three simulations with 6TIM crystal structure with position restraints on residues 1 to 150 (Pres-6TIM). We wanted to freeze motion of the first 150 residues which includes loop-5 (127 to 137). Loop-6 of chain-B in 6TIM is closed in the starting structure and during the simulation residues 172 to 177 adopt open conformation while N-terminus of loop-6 (167 to 169) is hindered by loop-5 and stays closed (**Figure 12**). From our Pres-6TIM simulations it can be seen that change in conformation of loop-5 from closed to open is required for loop-6 N-terminus to adopt open conformation. But in chain-A where loop-5 is arrested in open conformation, loop-6 N-terminus can sample closed conformation.

**Figure 12:**
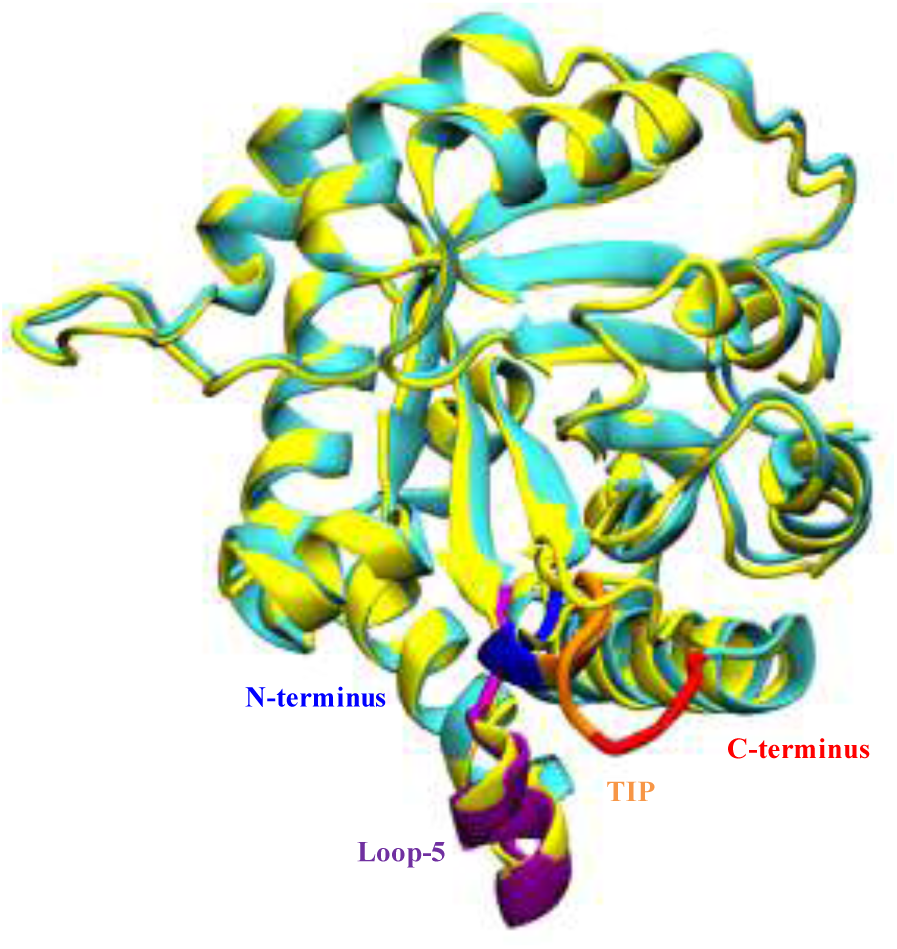
Snapshot of chain-B from the Pres-6TIM simulation where first 150 residues are position restrained in which N-terminus (blue) of loop-6 is closed because loop-5 is also in closed conformation. Solvent exposed tip (orange) and C-terminus (red) of loop-6 are open. Starting structure is in yellow.

#### 3.4.5 Loop-6 transition rate

Based on the W169-S212 and V176-S212 distances, average number of transitions in various apo and ligated ensembles including the unit cell simulations was calculated (**Figure 13**). A transition is counted as one jump from open to closed conformation of loop-6 and back to open state. It has to be noted we do not observe a transition in every trajectory between open and closed conformations of loop-6. In few trajectories we have as many as 20 transitions (**Figure 13**).

**Figure 13:**
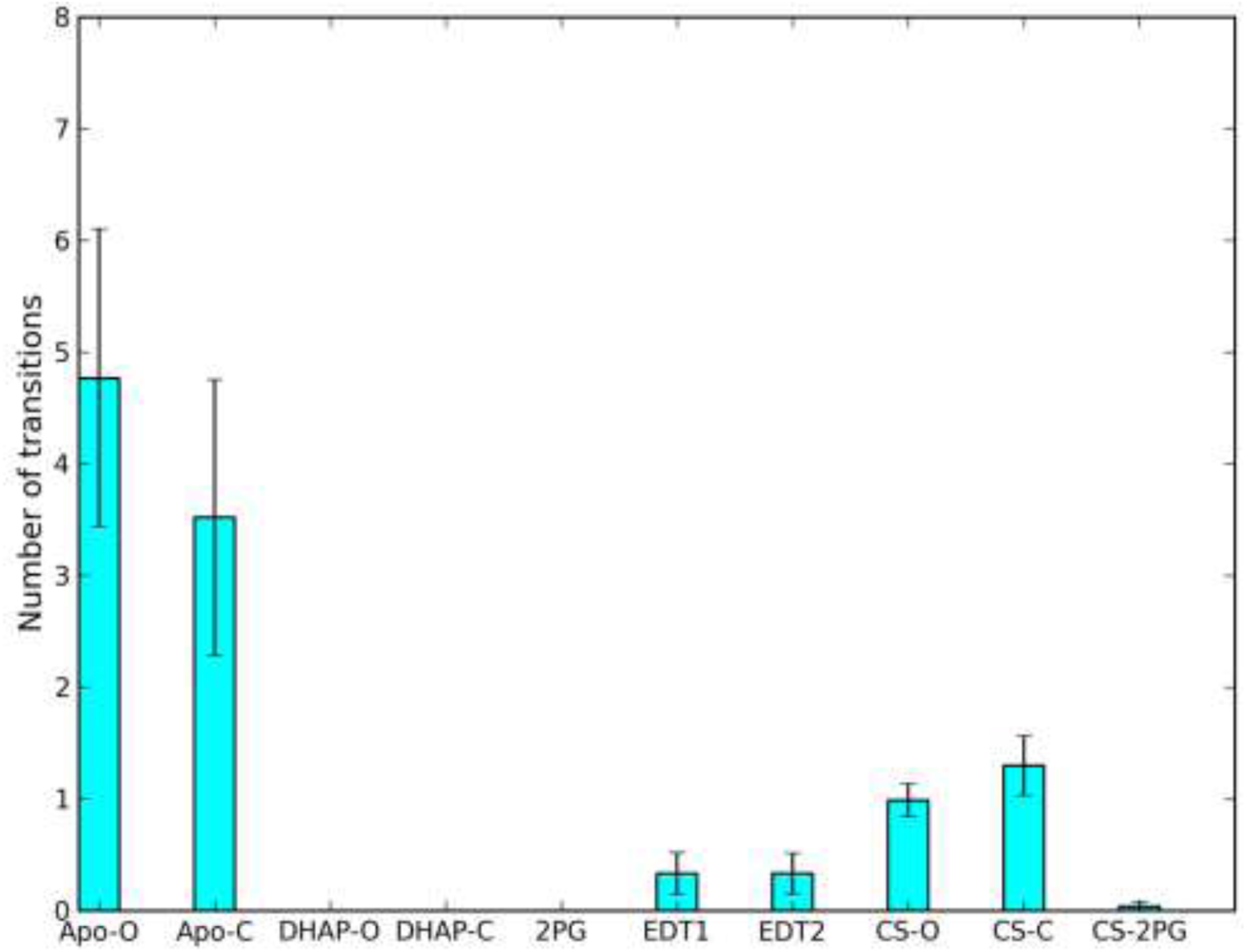
Average number of transitions of loop-6 between open and closed conformations. For ligated ensembles only that part of the trajectory during which ligand is bound in the active is considered.

The difference in number of transitions observed between various ensembles should be interpreted cautiously because our simulations have not converged i.e. in trajectories started with the same starting structure, in some trajectories we have seen transitions throughout the trajectory and in some simulations no transitions at all. Based on the transitions we have calculated the exchange rate between (k_ex_) open and closed conformations can be estimated which is on the order of 10^6^ s^−1^ .

In the ligated trajectories of DHAP-C and 2PG where we start with closed loop-6 and DHAP-O where we start with open loop-6, we have no transitions between open and closed conformations of loop-6 as long as the ligand is present in the active site. In DHAP-C and 2PG as stated earlier, loop-6 undergoes a conformational change to open conformation in the first 10 ns of the simulation. This must not be interpreted as, when the ligand is bound to the active site, loop-6 does not undergo a conformational change as we do see transitions in EDT1, EDT2 and CS-2PG. We need to perform longer simulations with ligand bound to the active site as we did for apo TIM simulations which were 1 μs long.

#### 3.4.6 Loop-6 principal component analysis

Loop-6 undergoes conformational change between open and closed states in apo, holo and unit cell ensembles on microsecond time scale. But as suggested by the X-ray data we do not observe a preference for closed loop-6 when ligand is bound to the active site. The experimental estimate for the free energy difference (ΔG) between open and closed states is given in **Table 1**. In apo TIM with open loop-6 as major conformation, the ΔG between open and closed conformations of loop-6 is 1.23 kcal.mol^−1^ to 1.77 kcal.mol^−1^. Based on the X-ray data when the ligand is bound to the active site it is expected that loop-6 is closed with ΔG of −1.23 kcal.mol^−1^ to −1.88 kcal.mol^−1^ .

**Table 1:** Estimates of the free energy difference between open and closed conformations of loop-6 based on population ratios from various NMR experiments.

| Apo/holo | Open ($p_a$) | Closed ($p_b$) | $\Delta G$ (kcal.mol<sup>-1</sup>) |
| --- | --- | --- | --- |
| **APO** | 8/20 | 1 | 1.23 to 1.77<sup>18</sup> |
| **2PG/G3P** | 1 | 8/20 | -1.23 to -1.77<sup>18</sup> |
| **APO (V167)** | 0.74 | 0.26 | 0.62<sup>22</sup> |
| **APO (T177)** | 0.91 | 0.09 | 1.37<sup>22</sup> |
| **G3P** | 0.04 | 0.96 | -1.88<sup>94</sup> |

To calculate the free energy difference between open and closed states of loop-6 from MD simulations first we have to design a reaction coordinate which describes this conformational change. Principal component analysis (PCA) is a very useful tool to identify, isolate and describe the large scale motions present in proteins. PCA was performed on chain-A of 5TIM (open loop-6) and chain-B of 6TIM (closed loop-6) X-ray structures, using the backbone atoms of the entire protein, excluding all the dimer interface loops (loops 1 to 4), loop-5, 7 and 8 along with N (residues 1 to 10) and C (241 to 249) terminus fragments. Eigenvector-1 (Ev-1) described the motion of loop-6 from open to closed conformation. All the TIM ensembles were projected along the Ev-1. In **Figure 14** to and fro sampling of open and closed states is seen along Ev-1. Using the probability distribution along Ev-1 we can estimate the ΔG between the open and closed states of loop-6 in various ensembles from equation 1.1.

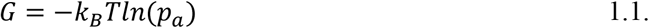

**Figure 14:**
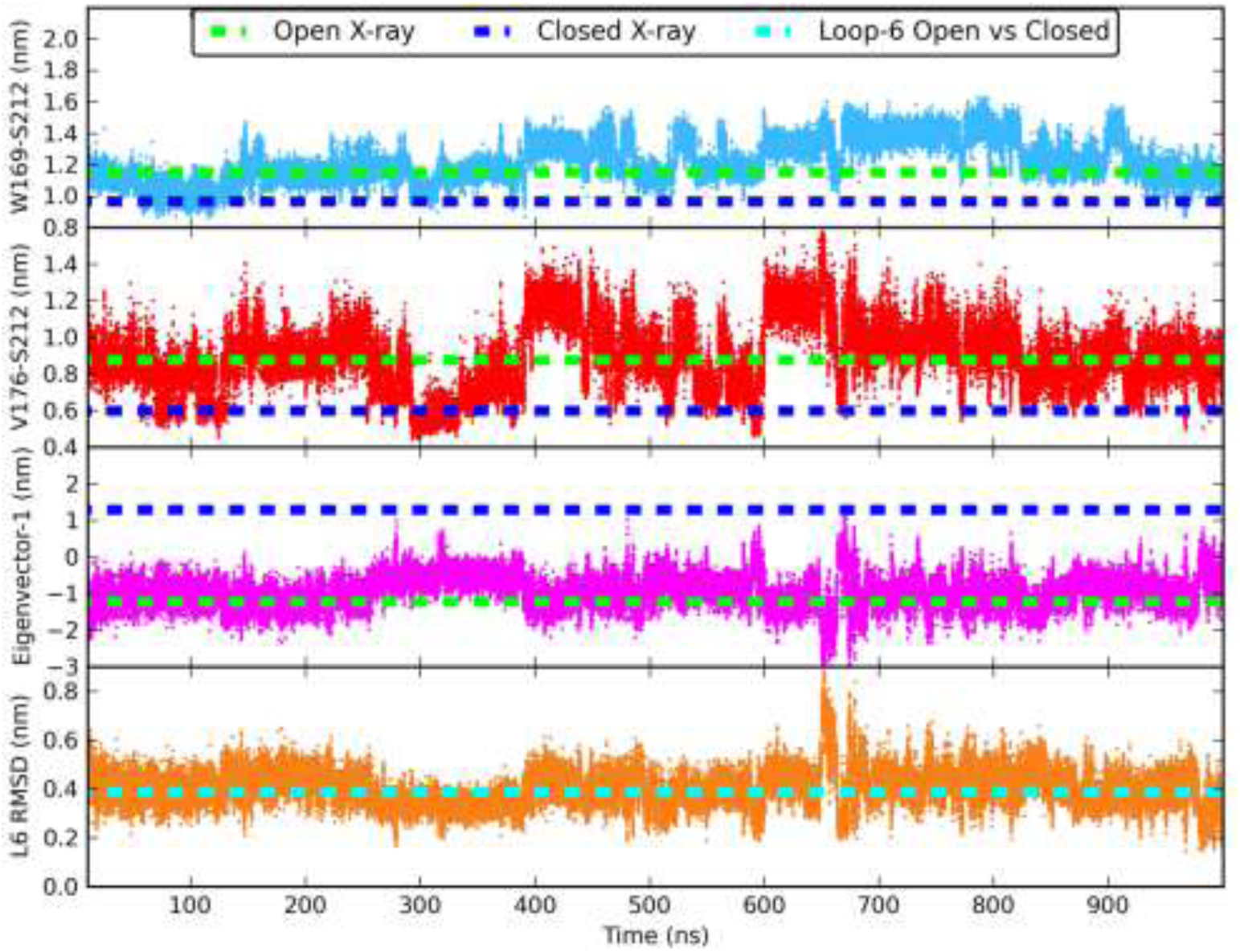
In panel-1 and panel-2 W169-S212 (blue), V176-S212 (red) distance evolution along time starting with a CTIM trajectory are shown. In the third panel we have plotted the projection of the same trajectory onto Eigenvector-1 (Ev-1, magenta) describing loop-6 motion. In the fourth panel we have the loop-6 (L6) RMSD showing that indeed when loop-6 is closed based on W169-S212, V176-S212 and Ev-1 of loop-6, rmsd is smaller than or equal to 0.2 nm.

We have combined both Apo-O and Apo-C ensembles to create APO ensemble because we observe similar behavior of loop-6 in both the ensembles. We have done the same for DHAP-O and DHAP-C ensembles. The free energy along the Ev-1 was calculated using the entire ensemble data. The error in free energy was calculated by estimating the free energy from each trajectory of the ensemble and bootstrapping the obtained free energy values 10000 times. The height of the error bars reflects the measure of sampling in MD simulation at that particular point along the Ev-1. As more trajectories sample a particular point, the height of the error bars become smaller and regions where sampling is not sufficient we have larger error bars.

In APO TIM ensemble we observe two minima, the first minimum at OTIM X-ray structure (−1.2 nm, green dotted line) between −0.6 nm and −1.4 nm (**Figure 15**). A second shallow minimum is observed at the CTIM X-ray structure (1.3 nm, blue dotted line) between 0.9 nm and 1.5 nm. The free energy difference between open and closed loop-6 in APO TIM ensembles is 1.35±0.36 kcal.mol^−1^ which is in the range of experimentally predicted values of 1.23 kcal.mol^−1^ and 1.77 kcal.mol^−1^ (**Table 1**), suggesting that the population distribution of open and closed loop-6 structures is not different from population distribution in NMR experiments^18,20,22^ .

**Figure 15:**
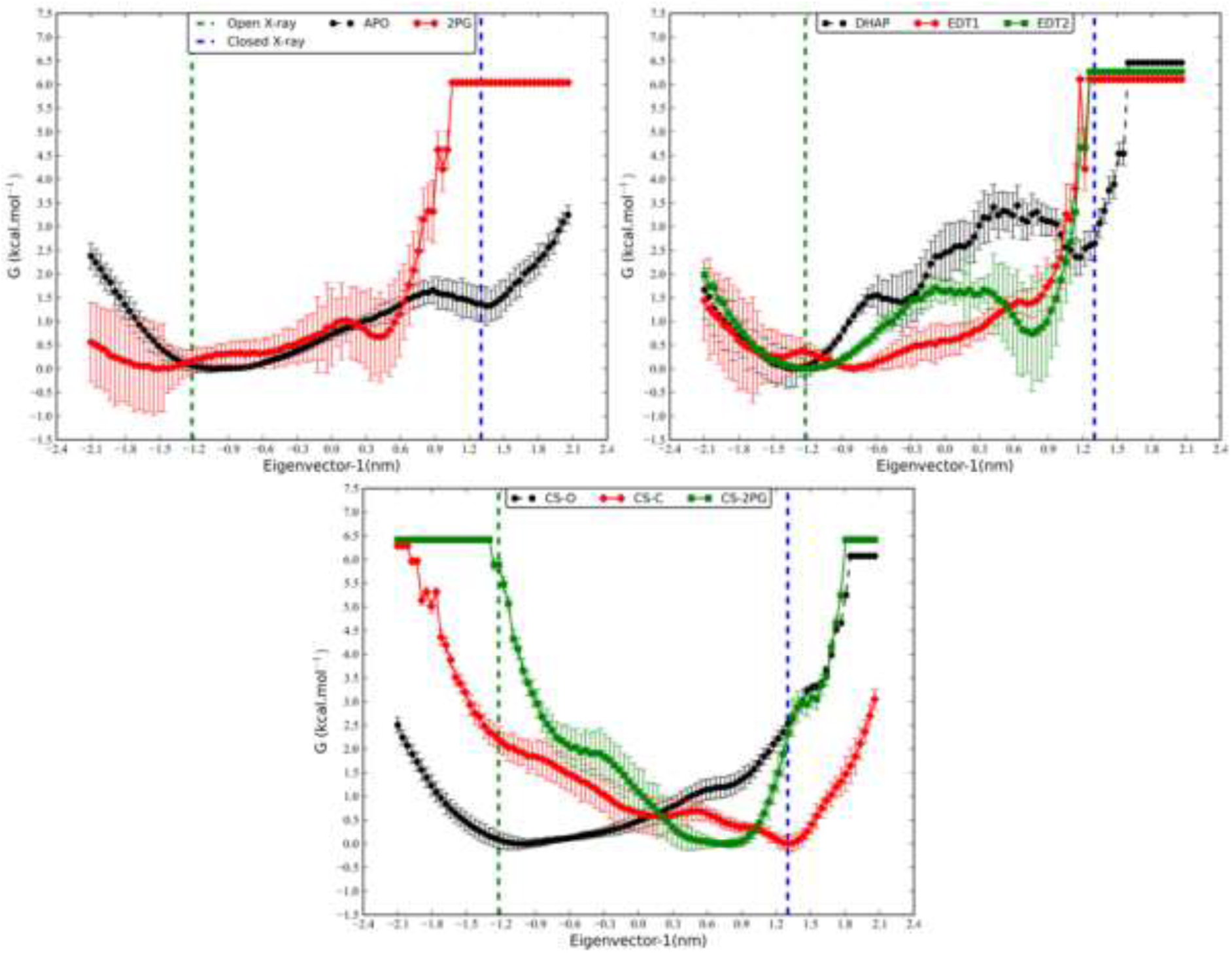
Loop-6 free energy profiles from various ensembles; apo, DHAP, EDT1, EDT2, 2PG, apo unit cell simulations started with open structure (CS-O), started with closed structure (CS-C) and ligated 2PG unit cell simulation ensembles.

ΔG calculated from ligated ensembles of DHAP, 2PG, EDT1, EDT2 is between 3.14±0.38 kcal.mol^−1^ to 4.96±0.22 kcal.mol^−1^ (**Figure 14**). The difference in the ΔG calculated from holo TIM simulations and the NMR holo ensemble can arise because of sampling issue i.e. the 300 ns simulation time for each trajectory might not be long enough to observe the ligand induced closed conformation for loop-6.

The ΔG calculated from apo unit cell simulations of CS-O and CS-C which start with OTIM and CTIM structure is 2.4±0.2 kcal.mol^−1^ favoring open loop-6 and −2.1±0.33 kcal.mol^−1^ favoring closed loop-6 (**Figure 15**). This is a surprising result especially the ΔG from CS-C ensemble which suggests that in the X-ray crystal environment the structures that are trapped in closed conformation have to work against the free energy gradient to reach the open state. We do observe sampling at but not any minima at reference CTIM X-ray structure for CS-O ensemble and at the OTIM X-ray structure for CS-C ensemble.

CS-2PG simulations start with a CTIM structure with 2PG bound to the active site pocket. The minima for closed loop-6 conformation, is seen between 0.6 nm and 0.9 nm (**Figure 15c**). RMSD analysis of loop-6 of structures from 0.6 nm to 0.9 nm of eigenvector revealed that loop-6 conformation resembles the X-ray closed loop-6 conformation (**Figure 16**). In CS-2PG crystal unit simulations closed loop-6 conformation is favored by −5.24±0.14 kcal.mol^−1^ (**Figure 15c**).

**Figure 16:**
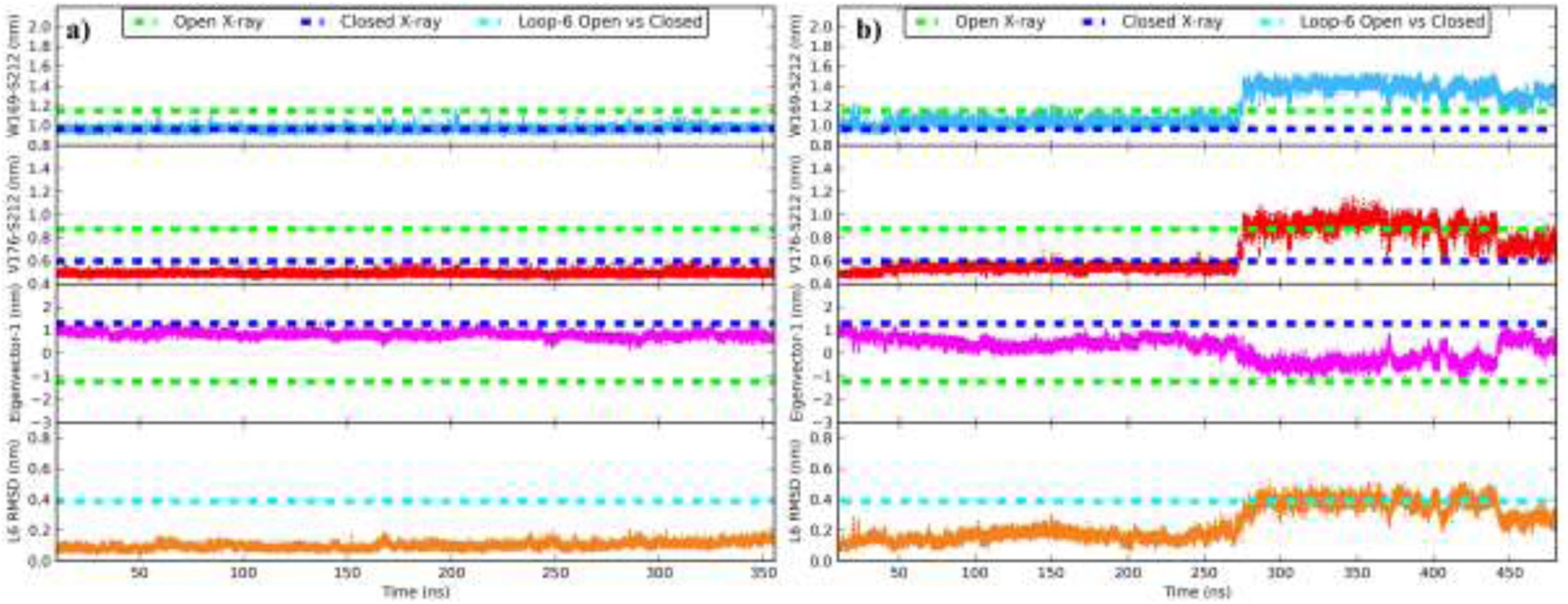
W169-S212 (blue), V176-S212 (red) distance evolution along time starting with a CTIM structure from CS-2PG ensemble trajectory. In the third panel we have plotted the projection of the same trajectory onto Eigenvector-1 (magenta) which describes loop-6 motion. In the fourth panel we have plotted the loop-6 (L6) RMSD (orange) showing that indeed when loop-6 is closed based on W169-S212, V176-S212 distances and projection of loop-6, rmsd is smaller than or equal to 0.2 nm and when loop-6 is open RMSD of loop-6 resembles X-ray open structure.

Williams et al.^18^ suggested that the ΔG between the 2PG ligated closed loop-6 and an unligated open conformation is −6.8 kcal.mol^−1^ if the binding free energy of 2PG (G_bind_ = −6.8 kcal.mol^−1^) is taken into account. Just based on loop-6 population ratios reported by Williams et al.^18^ and Massi et al.^94^ the ΔG in G3P/2PG ligated TIM would be between −1.23 kcal.mol^−1^ and −1.83 kcal.mol^−1^. To calculate the contribution of G_bind_ on ΔG we have to calculate the G_bind_ of each DHAP/EDT1/EDT2/2PG for open and closed loop-6 conformations, which would be the focus of future work.

#### 3.4.7 Free energy of Loop-6

The ΔG calculated from the apo TIM ensembles match the experimental values calculated from NMR population ratios. But the ΔG from holo TIM simulations of DHAP, EDT1, EDT2 and 2PG are in disagreement with the NMR data which might be because of sampling issue as suggested earlier in section 3.4.6. To overcome the sampling problem, we used enhanced sampling technique called Umbrella sampling (US).

We have demonstrated that loop-6 Ev-1 describes the transition of loop-6 between open and closed conformations (**Figure 13**). So it was chosen as the coordinate for performing umbrella sampling simulation (US) to calculate the free energy difference (ΔG) between open and closed conformations. The main reason behind excluding the dimer interface loops 1-4 and loop-5, loop-7 and loop-8 was the assumption that the regions of the protein which are correlated with loop-6 motion will adopt to the conformational change of loop-6 and also to avoid forcing of any region of protein along the reaction coordinate which does not contribute to this conformational change of loop-6.

Starting structures for each simulation were taken from eight individual trajectories with structures corresponding to eigenvalues along the reaction coordinate Ev-1. We performed totally eight US simulations, six apo (F5TIM-B, F6TIM-B, 5TIM-A, 5TIM-B, 6TIM-A and 6TIM-B) and two 2PG ligated simulations, 1N55-A and 1N55-B. We used 2PG in our US simulations over DHAP because of its longer residence time in the active site pocket and availability of NMR experimental data for comparison. As described in methods, umbrella windows were created between −1.60 nm and 1.60 nm with an interval of 0.2 nm. The OTIM X-ray structure projection on Ev-1 is at −1.22 nm and CTIM X-ray structure is at 1.29 nm.

All the trajectories from which structures were picked for US simulations, have sampled both open and closed states. If a particular, part along the reaction coordinate was not sampled in MD simulation, starting structure for that particular interval was created by linear interpolation. A harmonic restraining potential of 259 kcal.mol^−1^ was used for each umbrella window to restrict each structure at chosen point along the reaction coordinate.

The first 60 ns of each trajectory in US simulation was discarded as equilibration and only data from 60 ns to 110 ns was used to calculate the free energy profiles. US simulations from OTIM-B, CTIM-B, 6TIM-A, 6TIM-B, 5TIM-B and 1N55-A simulations will be referred to as loop-5 open US simulations (L5O) and 5TIM-A, 1N55-B as loop-5 closed US simulations (L5C). As demonstrated earlier in section 3.4.4, loop-5 motion is correlated with loop-6 N-terminus. A profound effect of loop-5 on loop-6 N-terminus conformation is seen in L5O US simulations. In L5C US simulations, in structures along the reaction coordinate from 0.6 nm to 1.4 nm where loop-6 is in closed conformation, loop-5 is also in closed state (**Figure 17a**). But in L5O US simulations, in the region between 0.6 nm to 1.4 nm, loop-5 adopts open conformation which causes loop-6 N-terminus to open (**Figure 17b**). This semi closed loop-6 where N-terminus is open because loop-5 is open, causes an increase in free energy difference between open and closed loop-6 states (Figure 17b).

**Figure 17:**
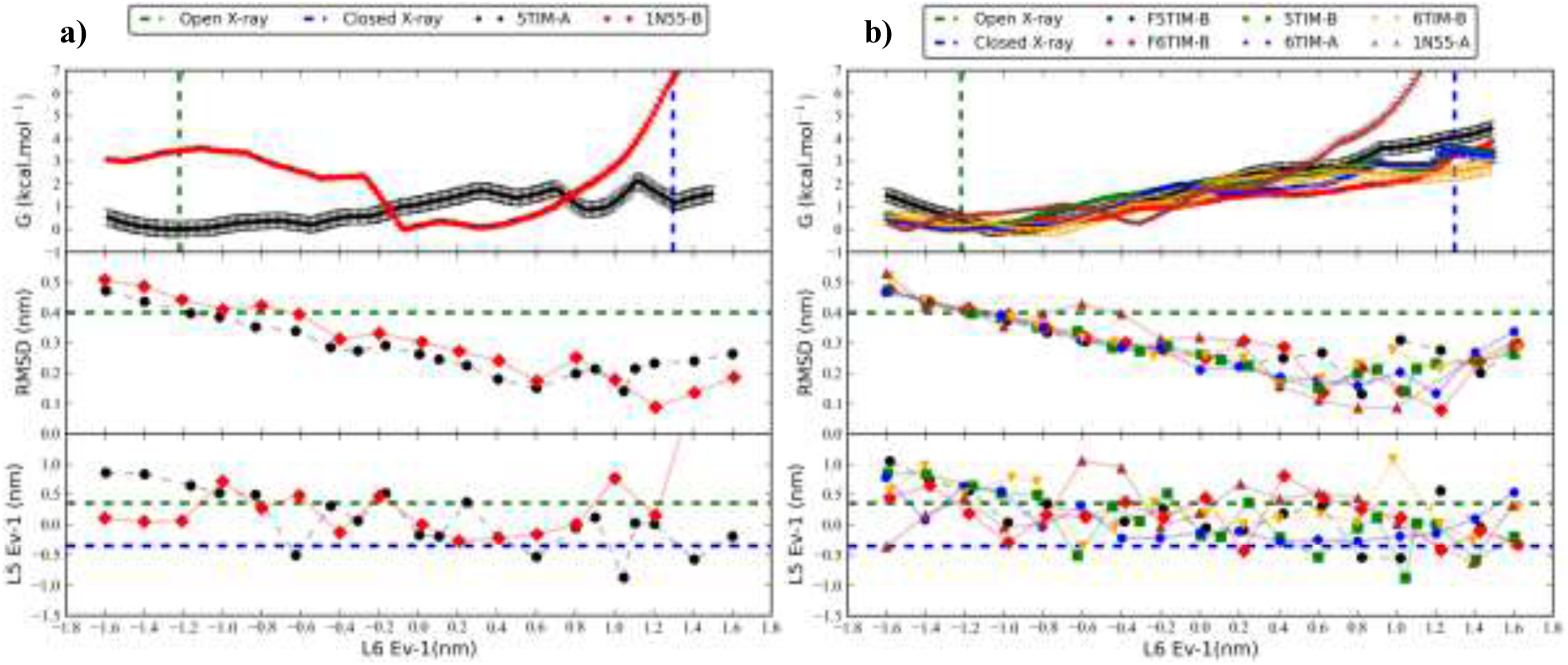
Free energy profiles from Umbrella sampling simulations from a) L5-Closed profiles and b) L5-Open profiles. In panel two and three each point represents the average of loop-6 RMSD with respect to the closed loop-6 (L6) in CTIM X-ray structure and loop-5 (L5) projection in eigenvector-1 (Ev-1) of loop-5 pca, which shows loop-5 conformational change between open (green) and closed (blue) loop-5.

The ΔG calculated from apo US simulation (ΔG_apo_) of 5TIM-A trajectory is 1.15±0.5 kcal.mol^−1^ (**Figure 17a**) which is comparable to the experimental value which lies between 1.23 kcalmol^−1^ and 1.77 kcal.mol^−1^ (**Table 1**). Whereas the ΔG calculated from 2PG ligated 1N55-B US simulation (ΔG2_PG_) is −3.4±0.2 kcal.mol^−1^ (**Figure 17a**) is far from the experimental estimate of −6.8 kcal.mol^−1^ and requires longer simulation time for each umbrella. Since we find profound influence of loop-5 on conformation of N-terminus of loop-6, a new reaction coordinate which includes loop-5 as well is required.

#### 3.4.8 Comparison with NMR

We have demonstrated that loop-6 transiently samples both open and closed conformations of loop-6 in apo and holo TIM ensembles. We have also defined loop-6 into five conformations and it is essential to segregate these conformations into NMR-Open (NMR-O) and NMR-Closed (NMR-C) since in NMR experiment only open and closed conformations are identifiable. We calculated the angular hop of W169 indole ring with respect to Z-axis from our MD simulations to compare with NMR data.

Ann McDermott and co-workers^18,20^ used W169 as an indicator of loop-6 conformational change. The angle between the W169 indole ring and the Z-axis differs by ~30° between OTIM (145°) and CTIM (114°) structures. We have calculated the angle between W169 indole ring and the Z-axis of the simulation box for every structure in the MD simulation. Based on loop-6 W169-S212 and V176-S212 distances, we have previously divided loop-6 into five conformations (**Supplementary Table 7**). We calculated the distribution of W169 angle for each conformation of loop-6 in all ensembles (**Figure 18**).

**Figure 18:**
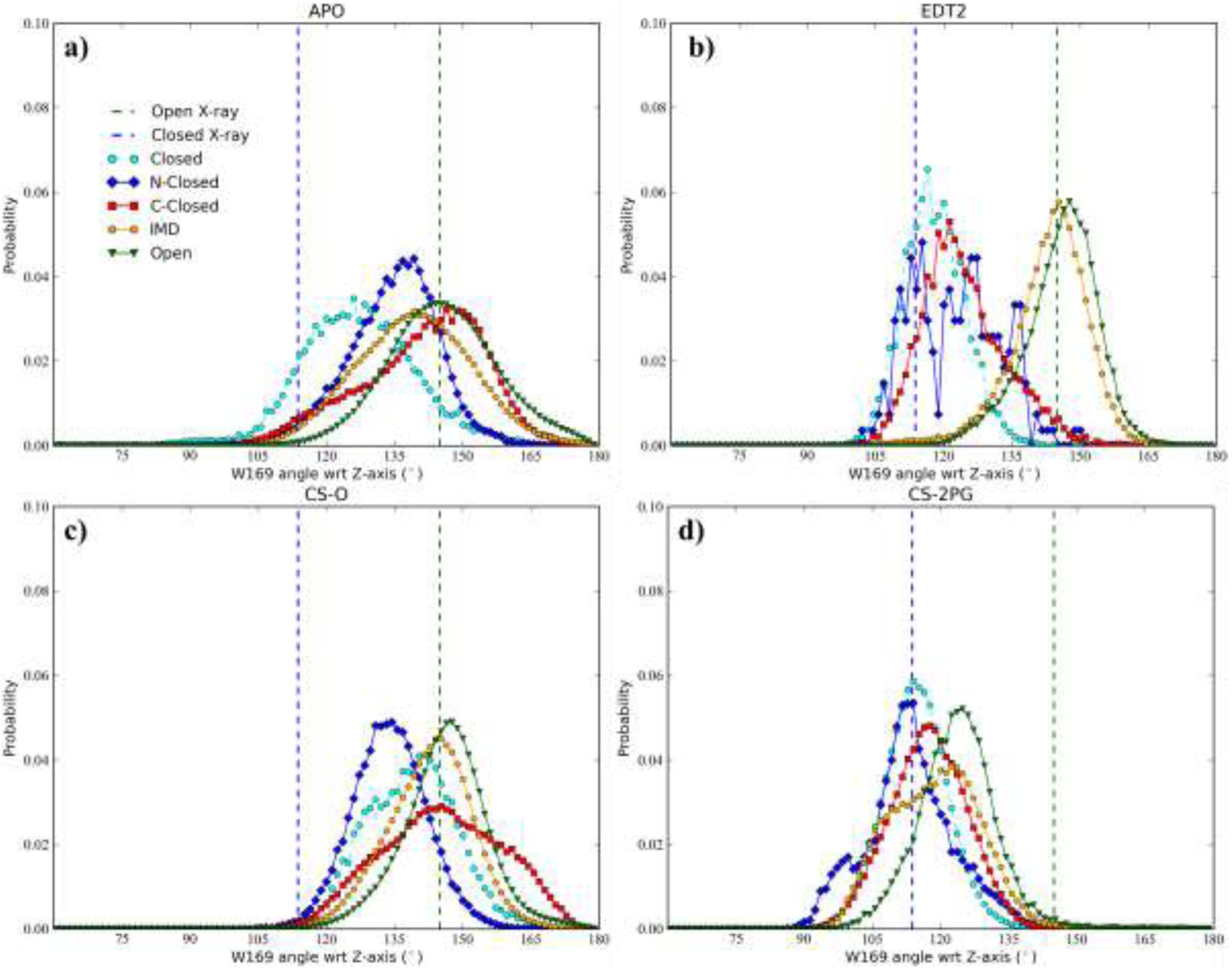
Distribution of W169 angle with respect to Z-axis for each of the five conformations of loop-6 in apo TIM ensemble (a), EDT2 ensemble (b), apo open unit cell simulation CS-O (c) and 2PG ligated unit cell simulation (d). When both termini of loop-6 are closed (Closed) or N-termini is closed (N-Closed) or C-termini is closed (C-Closed) we observe that W169 angle distribution shifts towards CTIM X-ray W169 angle. W169 angle distribution for loop-6 intermeditae conformation (IMD) and open conformation correspond to OTIM W169 angle.

For Open and IMD (loop-6 is neither closed nor open; **Supplementary Table 7**) loop-6 conformations, W169 angle corresponds to open W169 angle of 145°, which is seen in OTIM X-ray structure. When N-terminus of loop-6 is closed or both termini of loop-6 are closed we observe that W169 angle corresponds to CTIM X-ray structure value of 114° (**Figure 18**). W169 is present in N-terminus of loop-6 and it was expected that W169 would be only a reporter of closure of loop-6 N-terminus and entire loop-6. Surprisingly in crystal unit cell simulations and EDT1, EDT2 ligated TIM simulations, W169 angle corresponds to closed state even when C-terminus of loop-6 is closed and N-terminus is open. So we have combined Open, IMD as NMR-Open loop-6 states and C-terminus closed, N-terminus closed and loop-6 closed as NMR-Closed loop-6 states.

For each NMR-O and NMR-C conformation from MD ensembles, we calculated the average W169 angle with respect to Z-axis (**Figure 19**). In our simulations, in both apo and holo TIM ensembles the difference in W169 angle (ΔW169A) between NMR-O and NMR-C conformations is 6° to 10° with k_ex_ of 10^6^ s^−1^. The ΔW169A from apo TIM ensembles matches the SSNMR data from Rozovsky et al.^20^ in 2001.

**Figure 19:**
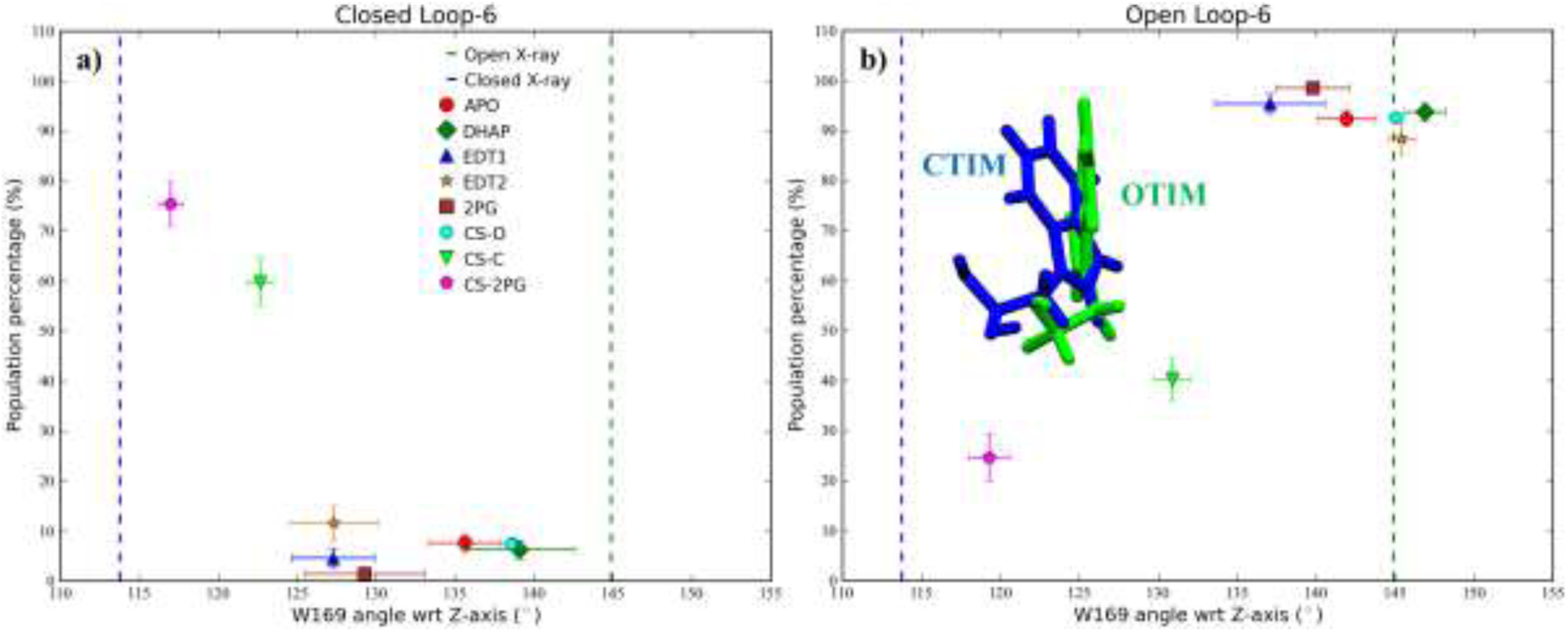
Average W169 angle with respect to Z-axis. On X-axis is the loop-6 population percentage in closed state (a) and open state (b).

Rozovsky et al.^20^ modeled the loop-6 conformational change in apo TIM sample with a low angular motion of 2° for W169 and k_ex_ of 10^6^ s^−1^ and in G3P ligated sample with a larger W169 angular motion of 20° to 40° with k_ex_ of 10^4^ s^−1^. Since we do not observe a preference for closed loop-6 in ligated TIM structures, the ΔW169A from holo TIM ensembles does not match the expected value reported by Rozovsky et al.^20^.

Williams et al.^18^ proposed a population ratio in the range of 1:8 to 1:20 for minor and major conformations with ΔW169A of 20° to 50°. In MD ensembles the population of the major conformation NMR-O is ~90% and of the minor conformation NMR-C is less than 10% except in CS-2PG (NMR-C 75%, NMR-O 25%) and CS-C (NMR-C 60%, NMR-O 40%) ensembles.

If we consider the ensembles with NMR-O/NMR-C populations of greater than 75% the ΔW169A value is 26° (**Figure 19**). This would imply that in SSNMR experiment if a sample has 100 OTIM structures, greater than 75% of the structures have to adopt closed loop-6 to observe a change in the line shape. We observe this in CS-C ensemble which starts with closed loop-6, 40% of the ensemble adopts NMR-O and 60% NMR-C conformation with W169 angles of 130° and 122° (**Figure 19**) and we would expect that as the population of NMR-O increases W169 angle would reach 145°. In apo TIM ensembles we do not see such a shift in population of loop-6 conformation.

### 3.5 Dynamics of loop-7

Loop-7 is found in open conformation in apo TIM structures and closed conformation is adopted in presence of the ligand molecules (**Figure 20**). Among all the apo TIM X-ray structures, loop-7 adopts closed conformation only in chain-B of 1R2R crystal structure. A water molecule is present in place of the oxygen of the phosphate moiety of the ligand, stabilizing the closed conformation of G211Ψ and S212Φ in the chain-B of 1R2R (**Figure 20b**).

**Figure 20:**
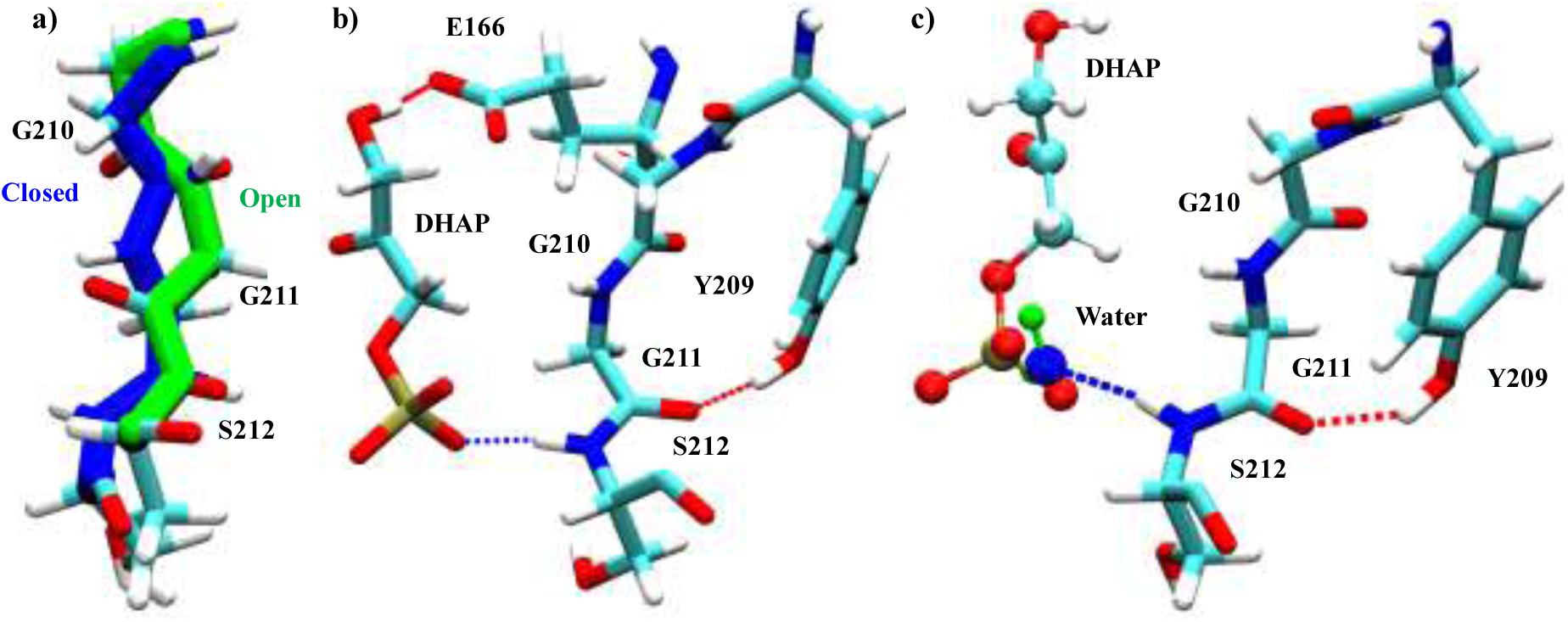
a) Open (green) and closed (blue) conformations of Loop-7. b) Loop-7 in closed conformation with DHAP bound to the active site. Hydrogen bond between amide group of S212 and phosphate moiety of DHAP along with hydrogen bond between the hydroxyl group of Y209 and carbonyl oxygen of G211 are only possible when G211Ψ and S212Φ dihedral angles flip to closed state. c) In chain-B of 1R2R crystal structure a water molecule (blue oxygen and green hydrogens) is present in place of the oxygen of the phosphate moeity of DHAP. This water stabilizes closed conformation of S212Φ. DHAP is shown in superposition with the water molecule.

We have performed multiple apo TIM and holo TIM simulations both in normal solvent and crystal unit cell simulations. We monitored the dihedral angles G210Ψ, G211Φ, G211Ψ, S212Φ, S212Ψ and V212Φ in various apo and holo TIM ensembles. G210Ψ, G211Φ, S212Ψ and V212Φ sample open and closed conformations in apo and holo TIM simulations (**Figure 21**). Whereas G211Ψ and S212Φ sample closed conformation only in presence of a water molecule located at a specific position or when the ligand molecules are bound to the active site (**Figure 21b**). In this work we would like to mainly focus on the dynamics of G210Ψ and G211Φ and its effect on conformation of catalytic residue E166 which is discussed in section 3.5.1 and in section 3.5.2 we discuss the dynamics of dihedral angles G211Ψ and S212Φ.

**Figure 21:**
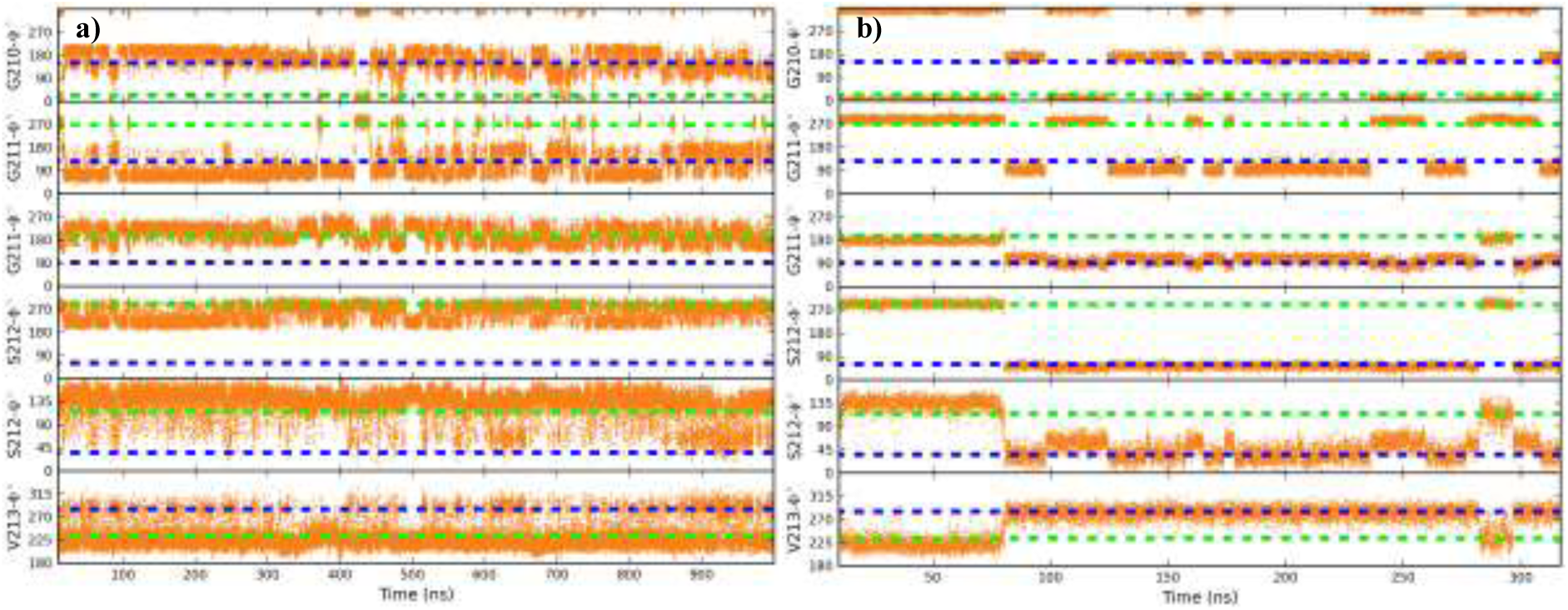
Dynamics of G210Ψ, G211Φ, G211Ψ, S212Φ, S212Ψ and V212Φ dihedral angles in an apo OTIM simulation (a) and DHAP ligated OTIM simulation (b). In both the simulations all the loop-7 dihedral angless start in open conformation. Apart from G211Ψ and S212Φ rest of the dihedral angles sample open and closed conformations in both apo and holo TIM simulations. Closed conformation of G211Ψ and S212Φ is ligand induced and in apo TIM simulations they prefer open conformation. The reference values for open (green dotted line) and closed (blue dotted line) conformations for each dihedral angles.

#### 3.5.1 Dynamics of G210Ψ and G211Φ

In both apo and holo TIM ensembles no effect of any ligand molecule is seen on the dynamics of G210Ψ and G211Φ dihedral angles. Casteleijn et al.^76^ proposed that, on ligand binding the closed conformation of G210Ψ induces a change in conformation of catalytic residue E166 from non-reactive ‘swung-out’ to reactive ‘swung-in’ conformation (**Figure 22a**). The clash between the carbonyl oxygen of G210 and P167 side chain was proposed to initiate this conformational change.

**Figure 22:**
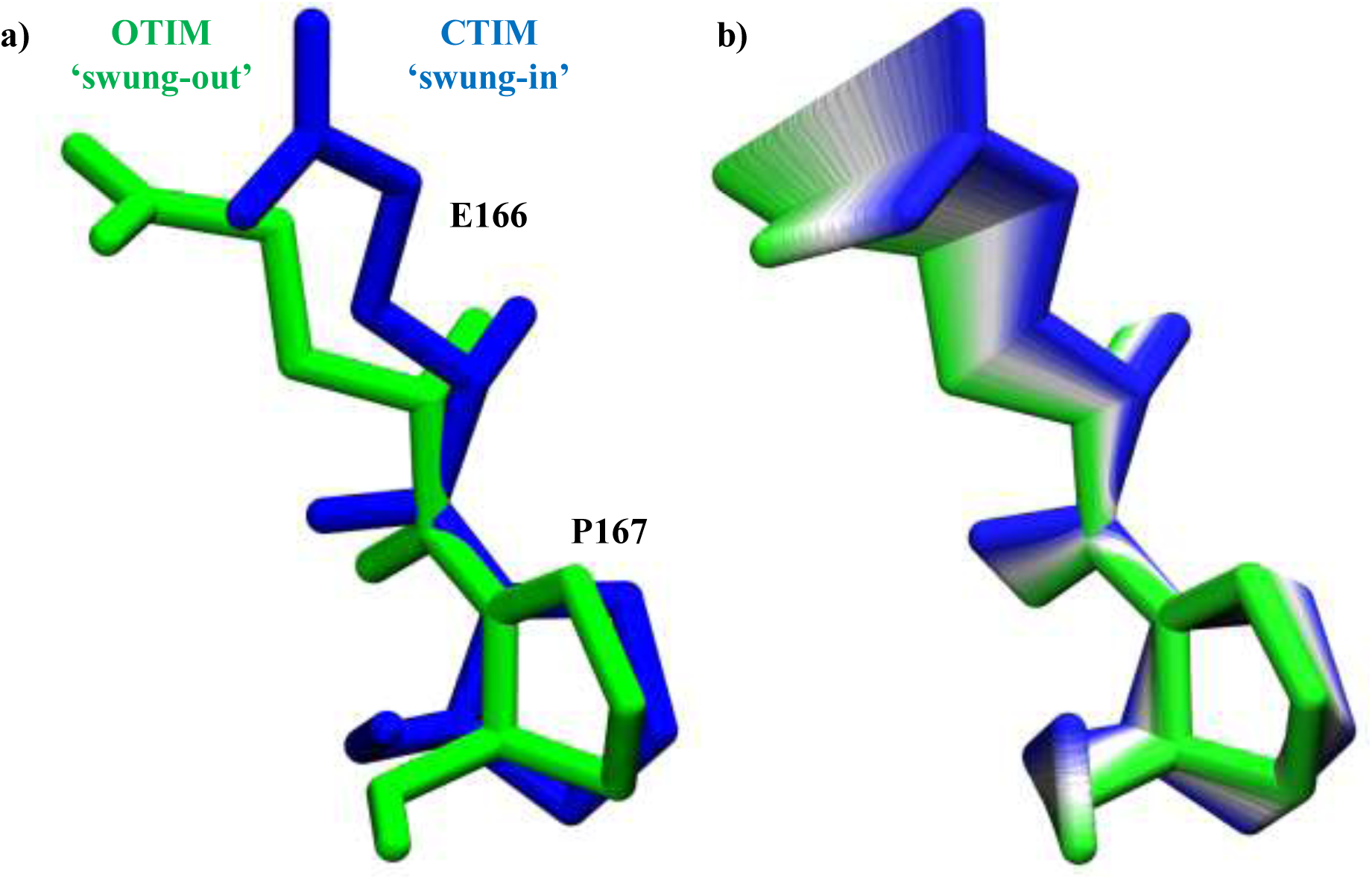
a) E166 swung-in (blue) conformation in CTIM and swung-out (green) conformation in OTIM. b) Conformational change from swung-in (blue) to swung-out oritentaion of E166-P167 along the first eigenvector from PCA.

To test this hypothesis, we have monitored the dihedral angles of G210Ψ and G211Φ along with the conformational change of E166. PCA was performed on the chain-A of 5TIM (open; **Figure 22a**) and chain-B of 6TIM (closed; **Figure 22a**). The first eigenvector (Ev-1) from the PCA described the conformational change of E166 from swung-in to swung-out conformation (**Figure 22b**).

We have monitored the dynamics of E166 along with the dihedral angles G210Ψ and G211Φ. G210Ψ and G211Φ are correlated i.e. when G210Ψ is in open conformation, G211Φ is also in open conformation and vice versa (**Figure 23**). Since E166 is present in the beginning of N-terminus of loop-6, we would expect that if E166 conformation is affected by loop-6 conformation then N-terminus of loop-6 will have the dominant effect. But we do not see such an effect of N-terminus on conformation of E166. E166 can sample both open and closed conformations independent of loop-6 N-terminus conformations (**Figure 23**) and same behaviour is seen across all TIM ensembles that we have simulated.

**Figure 23:**
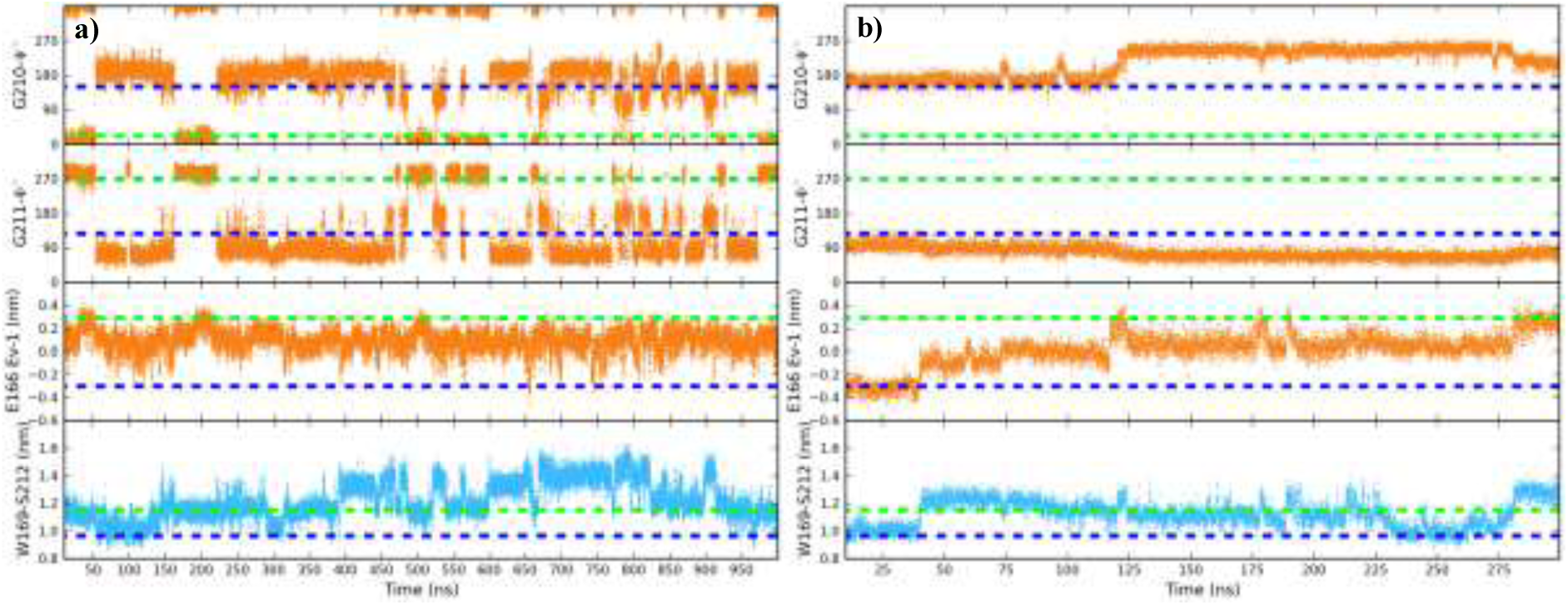
Dynamics of G210Ψ, G211Φ are shown in the first and the second panel. In the third panel, eigenvector-1 (Ev-1) from PCA of X-ray structures which describes the conformational change of E166 between CTIM swung-in conformation (dotted blue line) and OTIM swung-out conformation (green dotted line) is shown. Distance between C-alpha atoms of W169 present in N-terminus of loop-6 and loop-7 residue S212 is shown in fourth panel. In an apo CTIM simulation (a) and EDT1 ligated CTIM structure simulation (b) the change in conformation of G210Ψ and G21Φ also causes a change in E166 conformation along Ev-1.

E166 conformation is dependent on the conformations of dihedral angles G210Ψ and G211Φ. In simulations where G210Ψ and G211Φ stay in open conformation for the entire length of the simulation, E166 does not sample the swung-in conformation (data not shown). A deviation from the X-ray closed state for G210Ψ and G211Φ dihedral angles causes a change in the E166 conformation from swung-in to swung-out conformation (**Figure 23b**). To determine the orientation of G210Ψ and G211Φ dihedral angles when E166 is in swung-in or swung-out conformation, we calculated the average of G210Ψ and G211Φ dihedral angles for both swung-in and swung-out conformations of E166. A conservative cut-off of 0.3±0.3 nm for swung-out and −0.3±0.3 nm for swung-in was used to bin the dihedral angles assuming a two state population for E166 conformation.

If the conformation of dihedral angles G210Ψ and G211Φ induce the conformational change in E166 conformation as seen from our data, then we would expect that the average value of the dihedral angle over the entire ensemble when E166 is in swung-in conformation should correspond to the X-ray closed value (**Supplementary Table 2**) and when E166 is in swung-out conformation should correspond to X-ray open value (**Supplementary Table 2**). Since we observe that dihedral angles G210Ψ and G211Φ are correlated, here we only present the average values calculated for G210Ψ dihedral angle.

In **Figure 24** along x-axis we have shown the average value of G210Ψ angle and along y-axis the population percentage of E166 in swung-in (**Figure 22a**) or swung-out conformation (**Figure 22a**) in various TIM ensembles. As we expected above, we indeed observe that whenever E166 is in swung-in conformation G210Ψ dihedral angle corresponds to the X-ray closed state (**Figure 22a**) and when E166 is in swung-out conformation dihedral angle of G210Ψ samples the X-ray open state (**Figure 24b**). The large error bars for the population percentage represent the large variation in the population of E166 in swung-in or swuing-out conformation, suggesting that more sampling is required especially for the TIM ensemble with EDT1 bound to the active site pocket (**Figure 24b**). In CS-2PG simulation the percentage of swung-in conformation for E166 is ~3% which might account for the deviation of average of G210Ψ dihedral angle from X-ray open state (**Figure 24b**).

**Figure 24:**
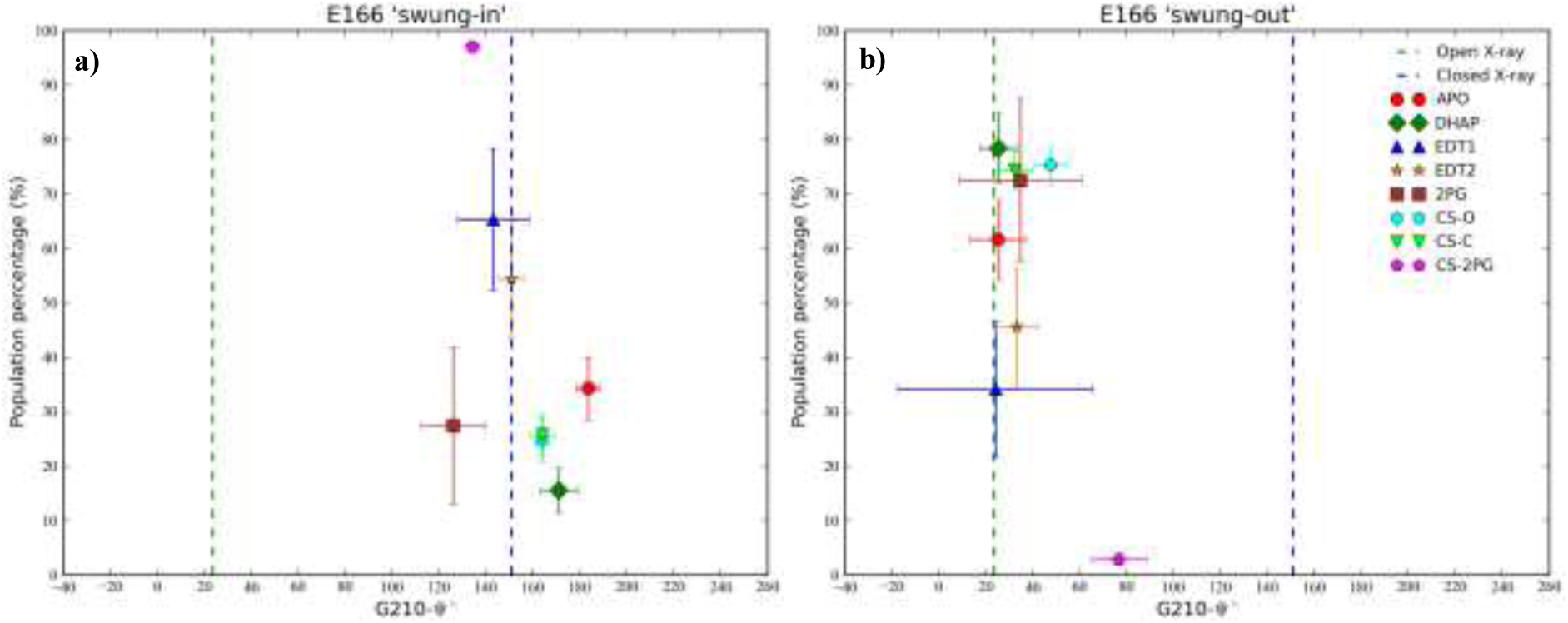
Average dihedral angle values of G210Ψ in various TIM ensembles when E166 is in swung-in conformation (a) and swung-out conformation (b). G210Ψ average open (green dotted line) and closed (blue dotted line) dihedral angle values from analyis of OTIM and CTIM X-ray structures (see section 3.1). Y-axis shows the population percentage of E166 in swung-in or swung-out conformations over the entire ensemble. Error bars were calculated from bootstrap analysis.

#### 3.5.2 Dynamics of G211Ψ and S212Φ

Dihedral angles G211Ψ and S212Φ sample the closed conformation only when the ligand molecules are bound to the active site or when a water molecule is present in place of the oxygen of the phosphate moiety of the substrate. In absence of this water molecule or the ligand, G211Ψ and S212Φ are present in open conformation. The effect of water molecules and ligand on the conformation of G211Ψ and S212Φ are further discussed in section 3.5.2.1 and 3.5.2.2.

##### 3.5.2.1 Effect of water

Apo OTIM and CTIM simulations were started using 5TIM and 6TIM X-ray structures. In apo OTIM simulations, out of 13 chains only in one chain G211Ψ and S212Φ flip from open to closed state (stays closed for 10 ns) and back to open conformation. In apo CTIM simulations, out of 25 chains, in five chains, G211Ψ and S212Φ transiently sample open and closed conformations. When G211Ψ and S212Φ flip from open to closed conformation, a water molecule similar to the one in the 1R2R crystal structure (**Figure 20b**) was found in these apo TIM simulations (**Figure 25**) which was not present in the beginning of the simulation. In rest of the apo OTIM and apo CTIM simulations, in absence of this water molecule at this particular position only open conformation is sampled by G211Ψ and S212Φ dihedral angles.

**Figure 25:**
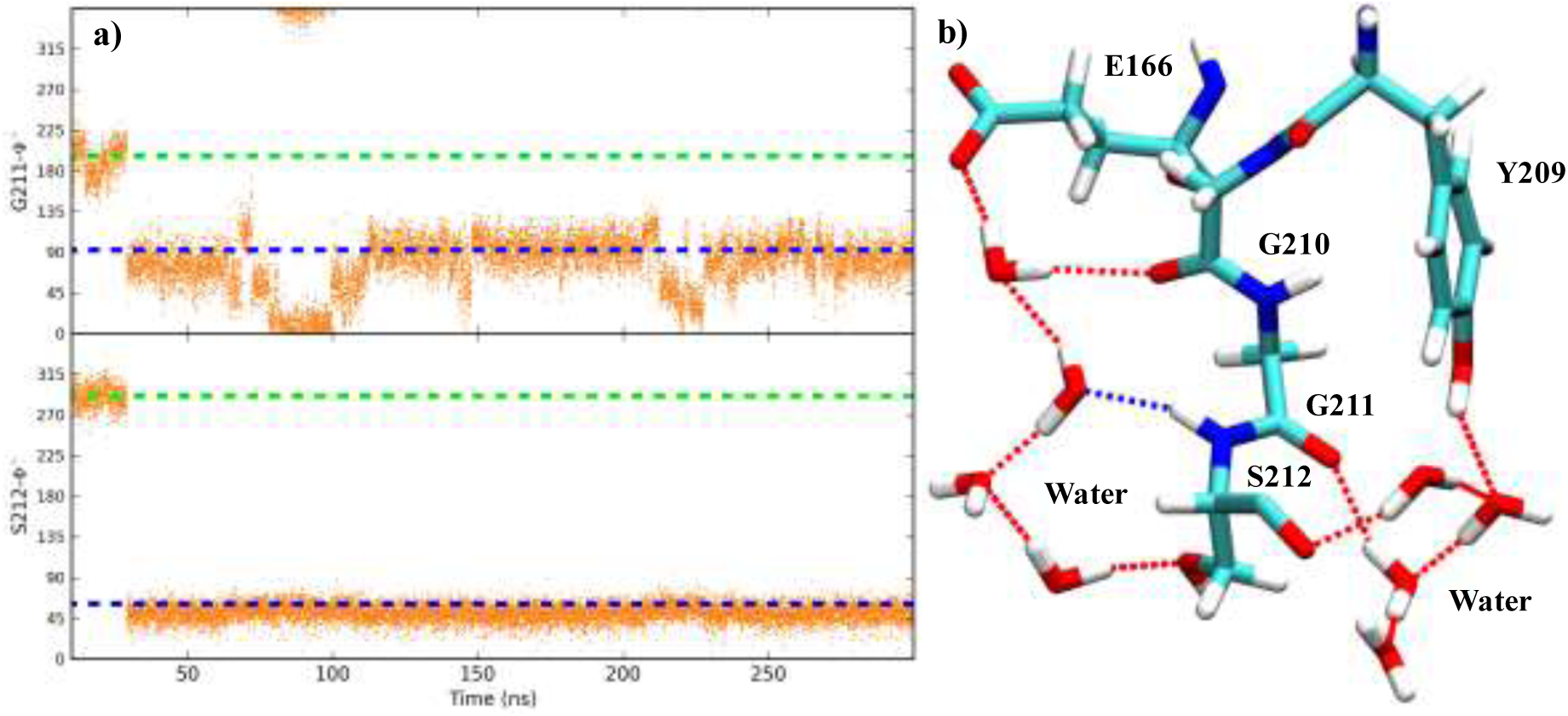
a) Apo crystal simulation where G211Ψ and S212Φ start in open conformation and flip to closed conformation. b) The closed conformation of G211Ψ and S212Φ in this apo crystal simulation is stabilized by water molecules.

The effect of water molecules on G211Ψ and S212Φ conformation is much more profound in apo crystal unit cell simulations which were started with 1R2R crystal structure. Out of 120 open chains, in 23 chains we observe that G211Ψ and S212Φ flip from open to closed state spontaneously. And in five chains G211Ψ and S212Φ flip to closed state and stay closed for 175 ns to 270 ns of simulation time which is not seen in the normal apo OTIM and CTIM MD simulations of 5TIM and 6TIM crystal structures. In these 23 chains, a water molecule which was not present in the beginning of the simulation was found within the hydrogen bonding distance of amide group of S212 stabilizing G211Ψ and S212Φ in closed state as seen in chain-B of the 1R2R X-ray structure (**Figure 20b**).

##### 3.5.2.2 Effect of ligand

As discussed in section 2.2 we performed multiple holo TIM simulations with natural substrate DHAP, inhibitor 2PG, reaction intermediates EDT1 and EDT2 in the active site pocket. DHAP was placed in the active site of both OTIM and CTIM structures whereas rest of the ligand molecules were placed in the active site pocket of CTIM structures. All the ligand molecules in the beginning of the simulation are bound to the active site in reactive complex (RC) orientation.

In section 3.4.1 we have shown that no influence of any ligand molecule is seen on the conformation of loop-6 i.e. even when the ligand molecules are bound to the active site pocket for entire length of simulation (~300 ns), loop-6 does not adopt the closed conformation. But in the holo TIM simulations which start with loop-7 in open conformation when DHAP is bound to the active site, dihedral angles G211Ψ and S212Φ flip from open to closed states spontaneously. And when the ligand escapes into the solvent within the first 40 ns from OTIM chains, such an effect is not observed. In section 3.1, we proposed that the flip of G211Ψ and S212Φ into closed state may be caused by the electrostatic repulsion between carbonyl oxygen of G211 and the negatively charged phosphate moiety of the substrate DHAP molecule.

We calculated the coulombic and Van der Waals interaction energy between the carbonyl oxygen of G211 and the amide group of S212 with phosphate moiety of natural substrate DHAP, reaction intermediates EDT1, EDT2 and the inhibitor molecule 2PG (**Figure 26**). The electrostatic repulsion between the negatively charged phosphate moiety and the carbonyl oxygen of the G211 can be as large is 4 kcal.mol^−1^ and this repulsion causes the rotation of G211Ψ (N-Cα-C-N). This rotation around G211Ψ simultaneously induces S212Φ into the closed conformation and brings amide group into the position to form hydrogen bond with phosphate moiety of the substrate. This flip to closed state provides additional −8 kcal.mol^−1^ of stabilization energy for the ligand molecule in the active site (**Figure 26a**).

**Figure 26:**
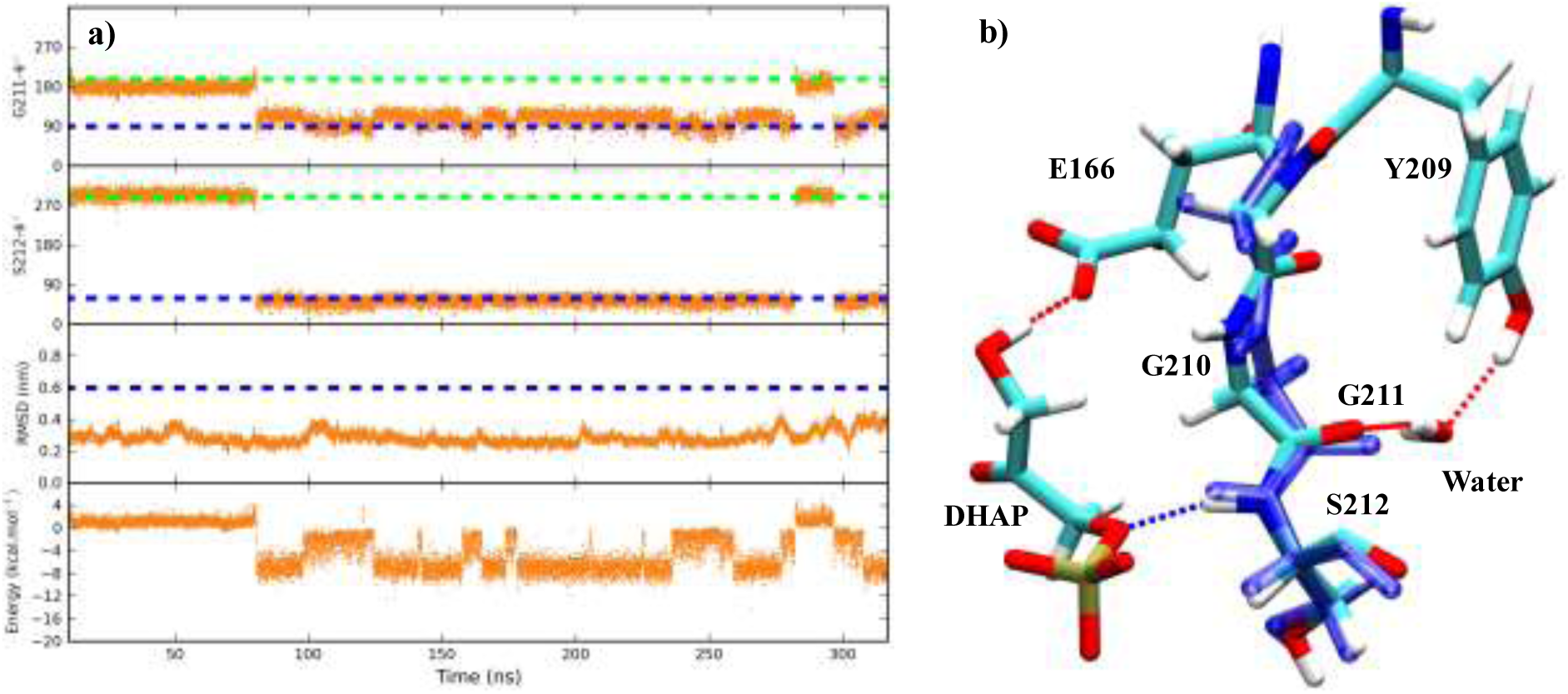
a) An OTIM structure which has DHAP bound in the active site pocket for the entire length of the simulation where G211Ψ and S212Φ flip from open to closed conformation. In the third panel RMSD of DHAP calculated with respect to the DHAP in the starting structure of the simulation.In the fourth panel the coulombic interaction energy between the phosphate moiety of DHAP and G211 carbonyl oxygen, S212 amide group is shown. b) In OTIM simulation loop-7 adopts a closed conformation in presence of DHAP. X-ray closed loop-7 is shown in transperent blue licorice. Dotted lines represent the hydrogen bonds.

The contribution of van der Waals interaction energy is compared to coulombic interaction energy between G211, S211 and phosphate moiety of ligand molecules is small. When G211Ψ and S212Φ are in open conformation because of the electrostatic repulsion between the phosphate moiety and carbonyl oxygen of G211, both the groups are apart which translates to a favorable Van der Waals interaction of less than −1 kcal.mol^−1^ . But when G211Ψ and S212Φ adopt closed conformation because of favorable electrostatic interactions (−8 kcal.mol^−1^) with phosphate moiety of ligand, steric clashes between phosphate oxygen and amide group of S212 increases Van der Waals energy up to ~3 kcal.mol^−1^ with average Van der Waals energy less than 1 kcal.mol^−1^ (data not shown). The free energy required to cause the flip of G211Ψ and S212Φ from open to closed state remains to be calculated.

## 4 Discussion

TIM is a highly efficient enzyme with k_cat_/k_m_ at 10^9^ M^−1^ s^−1^ and the reaction rate is only limited by diffusion of the substrate into the active site. To attain this catalytic efficiency, TIM has to streamline all the factors that contribute to the catalysis. These factors can be any conformational change involved in the catalysis, factors that affect the substrate binding and the product dissociation, as well as the barriers along the steps of the reaction mechanism.

In TIM structures, loop-6 and loop-7 envelope the active site and exist in open and closed conformations. Based on experimental studies it was proposed that the catalysis takes place when both loop-6 and loop-7 are in closed conformation. Loop-6 conformational change has been studied extensively studied by NMR experiments but not by microsecond MD simulations of entire TIM dimer. Dynamics of loop-7 conformational change is unexplored.

In this work we wanted to address the following questions. Is loop-6 and loop-7 conformation ligand gated or is it a natural motion of the protein? What factors affect the conformational changes in loop-6 and loop-7? What is the role of these conformational changes in the reaction catalyzed by TIM? How is the product formed at the end of the reaction released? Is there any influence of dynamics of loop-6 on loop-7 or vice versa?

### Loop-6 is not ligand gated but is influenced by loop-5

Loop-6 has a very important role to play in the reaction mechanism catalyzed by TIM. Based on X-ray structures loop-6 was proposed to exist in open and closed conformations. In apo TIM open loop-6 is the major conformation and in ligated TIM it is the closed loop-6. Closure of TIM loop-6 excludes water from the active site lowering the dielectric constant of which strengthens the electrostatic stabilization of the transition state^16^. Closure of loop-6 was also proposed to clamp the catalytic E166 between the two hydrophobic residues I171 and L231, which increases its basicity.

We find that as expected by Rozovsky et al.^19,20^, in apo TIM loop-6 does sample open and closed conformations at 10^6^ s^−1^ with W169 angular hop of 6°±2° (experimental value 2°). But when the apo TIM simulations (APO) start with a closed TIM structure (CTIM), a conformational change to open loop-6 is seen within the first 10 ns. Once the loop-6 in CTIM structures open up, for the rest of the simulation no difference is seen in loop-6 dynamics between simulations that start with open TIM (OTIM) structure and that start with CTIM structure.

In apo OTIM crystal unit cell simulations (CS-O) and in CTIM crystal unit cell simulations (CS-C), loop-6 like in APO ensemble samples open and closed loop-6 conformations at microsecond time scale. Unlike the normal MD simulations that start with CTIM structure, loop-6 stays closed for greater than 100 ns (Figured) with 60% of closed loop-6 conformation whereas the same in APO ensemble is 8%. As expected, this is because of slow solvent dynamics in the crystal unit cell simulations. On the other hand in CS-O percentage of open and closed loop-6 populations are 92% and 8% as seen in APO ensemble.

From Principal component analysis (PCA) using X-ray 5TIM and 6TIM structures we found eigenvector-1 (Ev-1) which described loop-6 conformational change. Using the distribution of projection along Ev-1, the free energy difference (ΔG) between open and closed loop-6 calculated from APO ensemble (1.35±0.36 kcal.mol^−1^) and apo loop-6 umbrella sampling (US) simulation (1.15±0.5 kcal.mol^−1^) is within the NMR experimental^18,22^ range of 1.23 kcal.mol^−1^ to 1.77 kcal.mol^−1^ which suggests that the population ratios of the major open and minor closed loop-6 conformation in apo TIM MD (9:1) and NMR (8:1 to 20:1) are identical.

The ΔG calculated from CS-O (2.4±0.2 kcal.mol^−1^) is higher than the NMR experimental^18^ value because of five times lesser transitions compared to the APO ensemble. Whereas the ΔG calculated from CS-C (−2.1±0.33 kcal.mol^−1^) favors closed loop-6 with a very strong minima observed at CTIM X-ray structure which implies that the crystal environment stabilizes closed loop-6.

The crystal unit simulations that start with apo 1R2R crystal structure from Rabbit at 100 K in which chains A, C and D are open and only chain-B is closed. The two other crystal structures 1R2T and 1R2S from the same organism but obtained at 85 K and 298 K do not show the same closed loop-6 conformation. Aparicio et al. pointed out to a water molecule which is present in place of the phosphate moiety of the ligand that might stabilize this closed loop-6 and further suggested that effect of flash freezing cannot be overlooked. Effects of flash freezing have been reported for side chain conformations of residues^88,90^ but not for such a large conformational change.

In the holo TIM simulations with natural substrate DHAP, inhibitor 2PG, reaction intermediates EDT1 and EDT2 we do not observe any conformational preference for closed loop-6 as expected from the X-ray data. The population ratio of open and closed loop-6 in normal holo TIM simulations is similar to apo TIM simulations with less than 10% of closed loop-6. Only in the 2PG ligated crystal unit cell simulation (CS-2PG) which starts with a CTIM structure, 75% of the ensemble has closed loop-6 and the rest 25% is open loop-6.

Based on NMR studies the expected k_ex_ between open and closed conformations is 10^4^ s^−1^ . This means that in the presence of the ligand, closure of loop-6 takes much longer time (once in 100 μs) than in the apo TIM structures (once in 1 μs). And our simulations with ligand are only 300 ns long. Though, in loop-7 we observe DHAP induced closed state (starting with an OTIM structure) in 50 ns of simulation time which is discussed in section 3.5. Such a spontaneous effect on conformation of loop-6 when ligand is bound to the active site, is not seen in our MD simulations. It is also possible that ligand induces conformational change from open to closed state in loop-7 first and loop-6 might follow later, which is not seen in our 300 ns long MD simulations.

The ΔG calculated from ligated ensembles is between 3.14±0.38 kcal.mol^−1^ and 4.96±0.22 kcal.mol^−1^ which is not close to the experimental value of −6.8 kcal.mol^−1^. Only the ΔG calculated from CS-2PG (−5.24±0.14 kcal.mol^−1^) is comparable to the experimental value. We performed two 2PG ligated US simulations 1N55-A and 1N55-B to calculate loop-6 free energy difference while 2PG is bound to the active site. The ΔG calculated from 1N55-A ensemble was ~2.7 kcal.mol^−1^ and from 1N55-B it was −3.4 kcal.mol^−1^ which is an improvement over normal MD simulation, but still not closed to the experimental value. The positive ΔG from 1N55-A is because of loop-5 open conformation in the CTIM structures along the reaction coordinate. A new reaction coordinate which also includes loop-5 conformational change is required to calculate more reliable estimates of ΔG from apo and holo TIM structures.

TIM catalyzes a reversible isomerization reaction where binding of DHAP results in formation of GAP and *vice versa*. A ligand induced conformational change for loop-6 as proposed by Richard and coworkers and X-ray data raises a very important question. How does TIM initiate opening of loop-6 once the reaction is completed? Donnini et al.^102^ has proposed a mechanism where closure of loop-6 stores energy as strain in ring of P167. Upon completion of reaction this energy is used to open loop-6. Such a mechanism also needs an instigator which signals completion of reaction, followed by opening of loop-6 and release of product.

A different proposition can be made where even when substrate molecules are bound to the active site, loop-6 undergoes a conformational change between open and closed conformations with the same k_ex_ of apo TIM (10^6^ s^−1^). In this scenario, when the substrate is bound to the active site, closure of loop-6 will enable the catalysis of the reaction. Since conformation of loop-6 is not ligand induced spontaneous opening of loop-6 will be followed by release of product and this will not require any trigger for initiating opening of loop-6, once it is closed. This proposal will be further discussed in section 5.3.

In our simulations we have monitored the dynamics of N-terminus (P167-V168-W169) and C-terminus hinges (K175-V176-A177) of loop-6 using distances between W169 and loop-7 residue S212 (W169-S212), V176 of loop-6 and S212 (V176-S212). Based on these W169-S212 and V176-S212 distances we divided loop-6 into five conformations. Not only do we observe different populations for the N and C-terminus hinges in open and closed conformations but also the correlation analysis between W169-S212 and V176-S212 distances has revealed that N and C-terminus hinges are uncorrelated i.e. loop-6 motion is not rigid.

N-terminus of loop-6 interacts with loop-5 whereas C-terminus of loop-6 is solvent exposed. For N-terminus of loop-6 to open loop-5 must also adopt open conformation. Hence higher correlation of 0.5 to 0.6 is seen between loop-5 and loop-6 N-terminus compared to the 0.2 seen between N-terminus and C-terminus distances. This effect of loop-5 is much more pronounced in US simulations.

The Ev-1 which was used as the reaction coordinate for US simulations only describes the loop-6 conformational change but not loop-5. The reason for leaving out loop-5 from Ev-1 was that if loop-5 conformational change is strongly correlated with loop-6, loop-5 will adopt to loop-6 conformational change. But in US simulations we observe that when loop-5 is in open conformation, loop-6 N-terminus hinge also adopts open conformation and does not stay closed. This suggests that loop-5 closure is essential to stabilize N-terminus of loop-6 in closed conformation.

To verify this, we have performed simulations with 6TIM X-ray structure where chain-B is closed and chain-A is open. The first 150 residues of the protein which includes loop-5 (residues 127 to 137) were position restrained. In these simulations, when loop-5 is restrained in closed conformation, loop-6 N-terminus stays closed but loop-6 C-terminus and tip undergo conformational change to open state. When loop-5 is restrained in open conformation in chain-A, both loop-6 N-terminus and C-terminus sample open, closed conformations.

We do not find any influence of any ligand molecule on conformation of loop-6 irrespective of its reactive complex (RC) or encounter complex (EC) orientation in the active site. From the analysis of X-ray structures we found out that when any ligand molecule is bound in EC state loop-6 is not present in closed conformation. In simulations with DHAP and CS-2PG when ligand is present in EC orientation closed conformation for loop-6 is observed. Phosphate moiety of loop-6 interacts with the tip of loop-6 (A170 to G174). We observe that closed conformation of N-terminus and C-terminus hinges is not hindered by phosphate moiety of ligand as is seen for tip of loop-6. When ligand is in EC orientation, tip of loop-6 opens in our simulations but N-terminus and C-terminus are not affected.

In all the OTIM X-ray structures, loop-5, 6 and 7 are in open conformation and in CTIM structure all the three loops are in closed conformation. In our apo TIM simulations when loop-7 (which will be discussed in 3.5) does not sample the closed conformation, loop-6 undergoes a conformational change between open and closed conformations. Because of this reason it might be possible that the closed conformation sampled in the NMR experiments (both loop-6 and loop-7 are closed) and MD simulations might be different. To confirm this, NMR studies which can study the conformational changes of loop-5, loop-6 and loop-7 are required.

### Loop-7 closure is induced by ligand/water

We have monitored the dynamics of dihedral angles G210Ψ, G211Φ, G211Ψ, S212Φ, S212Ψ and V212Φ in various apo and holo TIM ensembles. Based on X-ray crystal structures it was proposed that closed conformation of loop-7 is ligand induced^76,103^ . In our simulations we find that dihedral angles G210Ψ, G211Φ, S212Ψ and V212Φ undergo transitions between open and closed conformations in apo, holo and unit cell simulations and dihedral angles G211Ψ and S212Φ sample closed state only in presence of a ligand molecule or a strategically located water molecule as seen in chain-B of 1R2R crystal structure.

As proposed by Casteleijn et al.^76^ we also find that conformation of G210Ψ and G211Φ influence conformation of catalytic residue E166. Since we observe transitions between open and closed conformations for G210Ψ and G211Φ in both apo and holo TIM simulations, as suggested by Casteleijn et al.^76^ binding of ligand to the active site may not be necessary to induce closed conformation for G210Ψ and G211Φ. Depending on the conformation of G210Ψ and G211Φ, E166 undergoes conformational transitions between swung-in and swung-out conformations in apo, holo and unit cell simulations.

In the criss-cross pathway of the reaction mechanism, catalytic residue E166 removes the proton from C1 carbon of DHAP and moves to C2 carbon to donate the proton to the ketone oxygen^38,44^ . In this pathway E166 has to shuttle between the two carbon atoms of the substrate. Alahuhta et al.^104^ reported high mobility for the side chain of E166. In the TIM structure complexed with Phosphoglycolohydroxamate (PGH) which is a reaction intermediate analogue, E166 points away from the C2 carbon of the ligand (Supplementary Figure 2) whereas in TIM complex with 2-(N-Formyl-N-hydroxy)-amino-ethylphosphonate (IPP) which mimics the natural substrate GAP, E166 points at C2 atom of the ligand^104^.

If the flip of G210Ψ from open to closed conformation induces the *‘swung-in’* conformation, it would be an advantage to have the G210Ψ unperturbed by the presence of ligand in the active site. The change in this dihedral angle will aid in E166 movement between the two carbon atoms of the ligand.

Casteleijn et al.^76^ also suggested that when G210Ψ adopts closed conformation, the steric clashes between P167 and carbonyl oxygen of G210 might trigger closure of loop-6. In our apo TIM simulations even when G210Ψ stays in closed conformation, loop-6 conformation is not influenced i.e. loop-6 transiently samples both open and closed conformations.

Several mutations were created by Sampson et al.^25^ to probe the mechanism of closure of loop-6 on the active site of which Y209F had the most drastic effect with a 2400 fold decrease in the catalytic efficiency. Y209F mutation has minimal effect on the binding of the natural substrate GAP (less than 1 kcal.mol^−1^) and no formation of methylglyoxal is observed ^25^. Pompliano et al.^16^ created a TIM mutant where tip residues of loop-6 from I171-G172-T173-G174 were deleted which lead to the formation of methylglyoxal five times more frequently compared to wild type enzyme.

Sampson et al.^24^ measured the barriers for the reaction mechanism and found out the barriers for enolization increased by 5 kcal.mol^−1^ to 6 kcal.mol^−1^ in the Y209F mutant compared to the wild type. Based on the free energy profile Sampson et al.^24^ postulated that closure of loop-6 is affected because of loss of hydrogen bond between Y209 and A176 of loop-6 which in turn affects the reactive conformation of E166 accounting for the rise in barrier for proton transfer.

Before the work of Casteleijn et al.^76^ in 2006, it was speculated that closure of loop-6 induced conformational change in E166. But now it is clear that the change in dihedral angles of G210Ψ and G211Φ regulate the conformational change in E166 and conformation of loop-6 does not affect E166. Berlow et al.^22^ using NMR experiments in apo TIM has showed that Y209F mutation destabilizes the closed conformation by 0.84 kcal.mol^−1^, which does not account for the large change in the barrier of enolization reported by Sampson et al.^24^. Based on these new results, an alternative hypothesis can be postulated where Y209F mutation though it affects the stability of the closed loop-6 conformation, might have a larger affect on the conformation of G210Ψ and G211Φ dihedral angles in turn affecting the conformational sampling of E166. This hypothesis has to be tested by performing atomistic MD simulations of Y209F and comparing the populations of E166 in swung-in and swung-out conformation.

Our apo and holo TIM simulations have demonstrated that there is a pronounced influence of ligand and water molecules on the conformation of G211Ψ and S212Φ dihedral angles.Electrostatic repulsion of, 4 kcal.mol^−1^ between the negatively charged phosphate moiety of the ligand molecule and the carbonyl oxygen of G211 induces closed conformation for G211Ψ and S212Φ dihedral angles. The change in conformation of these two dihedral angles provides around −8 kcal.mol^−1^ of additional stabilization energy to the ligand molecule. As seen in chain-B of 1R2R crystal structure, in our apo open unit cell simulations we also found a water molecule which can stabilize the closed conformation of G211Ψ and S212Φ for 175 ns to 270 ns.

Amyes et al.^80^ reported that phosphate moiety of the substrate molecule induces closed conformation for loop-6 which we do not observe in our holo TIM simulations. Rather we suggest that phosphate moiety induces closed state for dihedral angles G211Ψ and S212Φ. To summarize there is no effect of ligand molecules DHAP, EDT1, EDT2 2PG on conformation of dihedral angles G210Ψ, G211Φ, S212Ψ and V212Φ whereas closed conformation of G211Ψ and S212Φ is ligand induced. As seen in 1R2R-B crystal structure, a water molecule can also induce closed conformation for G211Ψ and S212Φ. The free energy required for loop-7 conformational change between open and closed states, in presence of water and various ligand molecules has to be calculated in future.

To summarize, loop-7 might play a dual role in TIM’s catalysis. Dihedral angles G210Ψ and G211Φ are not affected by ligand’s presence in active site and might help shuttling of protons in the reaction mechanism by regulating the conformation of E166. Whereas closed conformation of G211Ψ and S212Φ is only sampled in presence of ligand molecule or a strategically placed water molecule. The closed conformation of G211Ψ and S212Φ removes the electrostatic repulsion between the carbonyl oxygen of G211 and the phosphate moiety of the ligand molecules and enables the formation of a hydrogen bond between the amide group of S212 and phosphate moiety’s oxygen, anchoring the ligand molecule in the active site.

## 5 Conclusions

Below, we have summarized the new insights into the dynamics of these two loops (loop-6 and 7) and loop-5 and then we discuss the plausible sequence of events in TIM’s catalysis.

### 5.1 Loop-6

We have shown that loop-6 spontaneously undergoes transitions between open and closed conformations in apo TIM structures at k_ex_ of 10^6^ s^−1^. As expected from the X-ray data, no preference for closed loop-6 is observed in our simulations with DHAP/EDT1/EDT2/2PG bound to the active site. Either the closure of loop-6 in presence of these ligand molecules is a much slower process than the simulation length of each trajectory, or loop-6 as in apo TIM simulations does not close even in the when one of these ligand molecules bind to the active site. We also find that closure of N and C-termini is not affected by ligand being in reactive conformation since it is the tip of loop-6 (170-AIGTG-174) that interacts with ligand and tip of loop-6 opens when ligand is in encounter complex orientation.

There is an effect of solvent dynamics on loop-6 conformational change. In the apo crystal unit cell simulations with 1R2R X-ray structure and 2PG ligated crystal unit cell simulations with 1N55 X-ray structure, because of ten times slower diffusion constant of solvent, opening times of loop-6 starting from closed conformation are ten times longer compared to the normal MD simulations. Nonetheless even in crystal the unit cell loop-6 transiently samples open and closed conformations.

Our simulations also strengthen the observation of Berlow et al.^22^ and Wang et al.^15^ that the population densities of loop-6 N-terminus and C-terminus in open and closed conformations are different. This suggests that loop-6 N and C-terminus do not move as a rigid body as proposed by earlier studies. The reason behind this is that, for N-terminus to open, loop-5 also has to adopt open conformation. Whereas C-terminus is solvent exposed and is not inhibited by any part of protein.

We find that, when loop-5 is restrained in closed conformation, loop-6 C-terminus samples open and closed conformations, whereas loop-6 N-terminus stays closed. When loop-5 is restrained in open conformation, loop-6 N and C-terminus sample both open and closed conformation. This confirms that loop-6 N and C-terminus are independent and that loop-5 conformation influences loop-6 N-terminus conformation. This uncoupled movement of N-terminus and C-terminus hinges of loop-6 and their influence on the reaction mechanism remains to be studied.

Effect of loop-5 conformation is also seen in loop-6 umbrella sampling (US) simulations. A restraining potential of 259 kcal.mol^−1^ was used to keep loop-6 closed along the reaction coordinate and when loop-5 adopts open conformation, on which there was no restraining potential applied, N-terminus of loop-6 opens. This further emphasizes importance of loop-5 in regulating conformation of loop-6 N-terminus. In future studies, to calculate loop-6 free energy difference between open and closed conformations, a reaction coordinate which describes both loop-5 and loop-6 conformational change from open to closed conformations should be used.

### 5.2 Loop-7

Loop-7 is a six-residue fragment (‘YGGSVN’) which also undergoes a conformational change. The backbone phi (Φ), psi (Ψ) dihedral angles of G210Ψ, G211Φ, G211Ψ, S212Φ, S212Ψ and V212Φ differ between open and closed conformations. Just like for loop-6, based on X-ray structures it was proposed that loop-7 closed conformation is ligand induced. With one exception i.e. in chain-B of 1R2R crystal structure both loop-6 and loop-7 are closed in the absence of a ligand. A water molecule is present in the active site in the place of a one of the oxygens of the phosphate moiety. This water molecule can form a hydrogen bond with the amide group of S212 stabilizing the closed conformation of loop-7.

In our simulations, we rather find that G210Ψ, G211Φ, S212Ψ and V212Φ sample open and closed conformations in both apo and holo TIM simulations. Whereas G211Ψ and S212Φ prefer open conformation in the absence of the ligand and adopt closed conformation only when a ligand molecule is bound to the active site. As seen in the chain-B of 1R2R crystal structure, we also find that in apo TIM simulations a water can also trigger closed conformation for G211Ψ and S212Φ. The role of S212Ψ and V212Φ is not known and was not studied in this work.

We monitored the conformational change of E166 between swung-in and swung-out conformations and G210Ψ between open and closed states. Our results suggest that G210Ψ conformation regulates the conformation of E166 in agreement with the proposal of Casteleijn et al.^76^. We propose that a flexible G210Ψ and G211Φ, which are not influenced by ligand, would be an advantage for the catalytic mechanism for shuttling protons between the carbon atoms by catalytic residue E166. This hypothesis has to be explored by further studies.

The change in conformation of G211Ψ and S212Φ leads to stabilization of the ligand in the active site. In apo TIM simulations G211Ψ and S212Φ prefer the open conformation and sample the closed conformation only when a water molecule is present within hydrogen bonding distance of S212 amide group. This water is similar to the water molecule seen in the chain-B of 1R2R crystal structure. In the absence of this water molecule G211Ψ and S212Φ never sample the closed conformation in apo TIM simulations. Energetics of this process, where water molecule induces closed conformation for G211Ψ and S212Φ is unclear and further studies are required.

We investigated the mechanism of G211Ψ and S212Φ conformational change in presence of a ligand molecule. When G211Ψ and S212Φ are in open conformation, the electrostatic repulsion between the carbonyl oxygen of G211 and the dianionic phosphate moiety of the ligand molecule induces a closed conformation for G211Ψ and S212Φ. In closed state as stated earlier, amide group of S212 and oxygen of the phosphate moiety can form a hydrogen bond and provide additional stabilization interactions to the substrate in the active site.

Amyes et al.^80^ suggested that some part of the phosphate binding free energy is used to induce closed conformation for loop-6. Based on our MD simulations we would rather suggest that it is rather utilized in inducing closed conformation for G211Ψ and S212Φ and not for loop-6 because even when ligand molecules (DHAP/EDT1/EDT2/2PG) are bound to the active site we do not observe closed conformation for loop-6. From our simulations it is quite evident that closed conformation of G211Ψ and S212Φ is induced by either by a ligand molecule or a strategically placed water molecule.

### 5.3 Plausible sequence of events

Base on previous studies, the existing view of TIM’s catalysis and sequence of events that enable the reaction are following. Binding of either of the substrates DHAP/GAP induces closure of loop-7. G210Ψ and G211Φ flip to closed state, triggers swung-in conformation for E166 and closed conformation for P167, which in turn closes loop-6. After the catalysis, loop-7 flips to open conformation followed by opening of loop-6 and release of the product.

From our results we propose the following scenario. Binding of ligand will induce a closed conformation for G211Ψ and S212Φ, which will an additional hydrogen bond to the substrate. Since the conformational change of G210Ψ is not influenced by binding of substrate, transitions between open and closed conformations will induce a conformational change in E166. This might help shuttling of protons by E166 between the two carbon atoms of the substrate.

Loop-6 transiently samples open and closed conformations. And when the substrate is bound to the active site, after closure of loop-6, catalysis will take place. If the closed conformation of loop-6 is not ligand induced, no special trigger is required to initiate opening of loop-6 and opening of loop-6 will provide an opportunity for release of the product.

Loop-6 has to stay closed long enough for catalysis to take place without the formation of methylglyoxal which is formed by solvent exposure of the reaction intermediates. It was suggested that the cellular environment where TIM performs this catalysis, mimics the crystal environment because of crowding in the cellular cytoplasm^105-107^. In our crystal simulations, starting from an open conformation, loop-6 undergoes a spontaneous conformational change to closed state, stays closed for greater than 100 ns and samples the open conformation again. Proton transfer occurs in sub picosecond time scales and since loop-6 stays closed for nanoseconds, this time duration should be sufficient for the catalytic process.

The catalytic rate of TIM is 10^4^ s^−1^ in GAP to DHAP direction and 10^3^ s^−1^ in GAP to DHAP direction, i.e. to catalyze one molecule of GAP to DHAP time required is 100 μs and for one molecule of DHAP to GAP it is 1 ms. We observe conformational changes in loop-6 and loop-7 dihedral angles G210Ψ, G211Φ, G211Ψ and S212Φ in microsecond regime. As long as the conformational changes that affect the catalysis are faster than or equal to the diffusion rate of the substrate, than the catalytic rate of the reaction is still limited by the diffusion of the substrate into the active site or product out of the active site.

